# Delineating antibody escape from Omicron sublineages

**DOI:** 10.1101/2022.04.06.487257

**Authors:** Henning Gruell, Kanika Vanshylla, Michael Korenkov, Pinkus Tober-Lau, Matthias Zehner, Friederike Münn, Hanna Janicki, Max Augustin, Philipp Schommers, Leif Erik Sander, Florian Kurth, Christoph Kreer, Florian Klein

## Abstract

SARS-CoV-2 neutralizing antibodies play a critical role in prevention and treatment of COVID-19 but are challenged by viral evolution and antibody evasion, exemplified by the highly resistant Omicron BA.1 sublineage.^1–12^ Importantly, the recently identified Omicron sublineages BA.2.12.1 and BA.4/5 with differing spike mutations are rapidly emerging in various countries. By determining polyclonal serum activity of 50 convalescent or vaccinated individuals against BA.1, BA.1.1, BA.2, BA.2.12.1, and BA.4/5, we reveal a further reduction of BA.4/5 susceptibility to vaccinee sera. Most notably, delineation of the sensitivity to an extended panel of 163 antibodies demonstrates pronounced antigenic differences of individual sublineages with distinct escape patterns and increased antibody resistance of BA.4/5 compared to the most prevalent BA.2 sublineage. These results suggest that the antigenic distance from BA.1 and the increased resistance compared to BA.2 may favor immune escape-mediated expansion of BA.4/5 after the first Omicron wave. Finally, while most monoclonal antibodies in clinical stages are inactive against all Omicron sublineages, we identify promising novel antibodies with high pan-Omicron neutralizing potency. Our study provides a detailed understanding of the antibody escape from the most recently emerging Omicron sublineages that can inform on effective strategies to prevent and treat COVID-19.

## Main

Despite increasing levels of immunity against SARS-CoV-2 induced by vaccination and infection, the Omicron variant resulted in a global surge of infections that reflects its high transmissibility and immune evasion mediated by the highly mutated spike protein.^1,4–15^ Although prolonged vaccine dosing intervals and booster immunizations based on the ancestral Wu01 strain elicit Omicron neutralizing serum activity, titers against Omicron remain considerably lower compared to those against other variants.^4,6,11,16–18^ Moreover, the spike mutations of Omicron have rendered several therapeutic monoclonal antibodies ineffective.^7,11,19^ Most experimental evidence on the resistance of Omicron to antibody-mediated neutralization is limited to analyses of the initial BA.1 strain. However, novel sublineages are rapidly emerging and associated with increasing numbers of infections.^20,21^ Determining their antibody escape properties is therefore of critical importance to effectively guide preventive and therapeutic measures. To this end, we analyzed in detail antibody-mediated neutralization of prevalent and emerging Omicron sublineages (BA.1, BA.1.1, BA.2, BA.2.12.1, and BA.4/5) both on a polyclonal and monoclonal level.

## Results

### Rapid spread of Omicron sublineages

The spike protein of BA.1 differs in 39 amino acid residues from the ancestral Wu01 strain of SARS-CoV-2 (**Fig. 1a**). While other Omicron sublineages share several mutations with BA.1, they diverge at various amino acid positions including in critical antibody epitopes (**Fig. 1a,b**). For example, the BA.1.1 spike protein contains an R346K substitution in the receptor-binding domain (RBD) that was also observed in the SARS-CoV-2 Mu variant and associated with escape from neutralizing antibodies (**Fig. 1a**).^22^ While BA.1 and BA.1.1 dominated the initial Omicron surge, they were rapidly outcompeted by BA.2 (**Fig. 1c**). BA.2 shares 21 of its 31 spike protein changes with BA.1 but differs considerably in both the N-terminal domain (NTD) and the RBD, regions targeted by the most potent SARS-CoV-2 neutralizing antibodies (**Fig. 1a,b**).

**Fig. 1.**
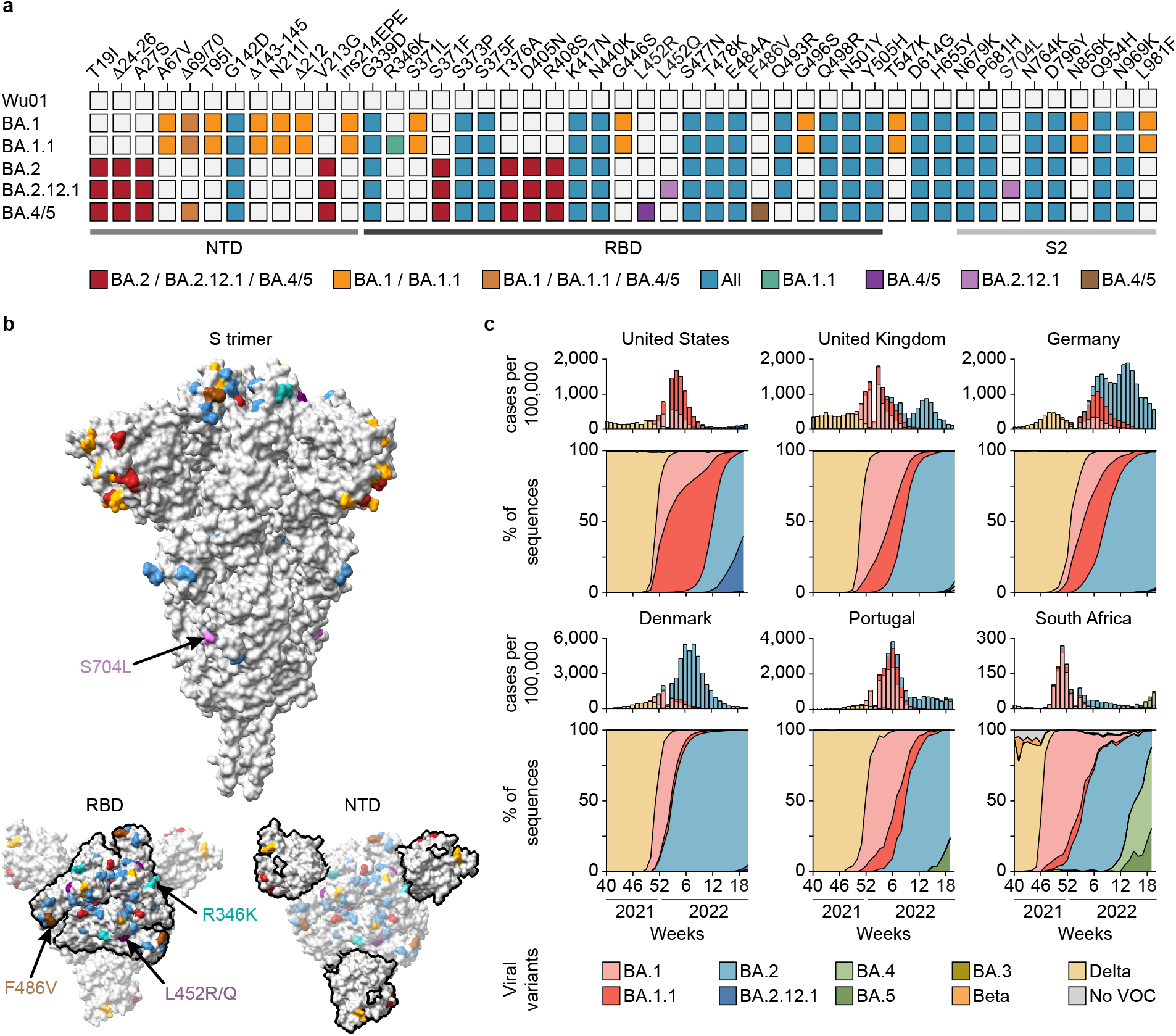
Omicron sublineage differences. **a,** Spike amino acid changes in the BA.1, BA.1.1, BA.2, BA.2.12.1, and BA.4/5 Omicron sublineages relative to Wu01. **b,** Locations of changed amino acids in Omicron sublineages highlighted on the SARS-CoV-2 spike (PDB: 6XR8), with colors indicating mutations shared between different sublineages. Representations at the bottom show RBD and NTD outlined in black. Changes exclusive to BA.1.1 are indicated in green and those to BA.4/5 in brown, residue 452 (mutated in BA.2.12.1 and BA.4/5) is indicated in purple. **c,** Top panels indicate weekly reported SARS-CoV-2 infections as aggregated by the COVID-19 Data Repository by the Center for Systems Science and Engineering at Johns Hopkins University and http://ourworldindata.org. Relative proportions of infections by variant per week were extrapolated from the proportion of sequences submitted to the GISAID SARS-CoV-2 database per variant and week (accessed on May 24, 2022) shown in the bottom panels. NTD, N-terminal domain; RBD, receptor-binding domain; PDB, Protein Data Bank.

Following a decline in cases from the BA.1-/BA.2-related first Omicron wave, new sublineages are rapidly emerging (**Fig. 1c**). For example, BA.2.12.1 outcompetes other BA.2 sublineages in the United States resulting in an increase of infections. Moreover, within weeks of their identification, the BA.4 and BA.5 sublineages with an identical spike protein have become dominant in South Africa and Portugal and driven rising case numbers (**Fig. 1c**).^21^ BA.2.12.1 and BA.4/5 share the majority of their spike mutations with BA.2 (94% and 90%, respectively), but contain additional changes at sites associated with antibody escape. These include substitutions at residue 452 (L452Q in BA.2.12.1 and L452R in BA.4/5, respectively) that were also recorded for the Lambda and Delta variants (**Fig. 1a**).^23,24^ In addition, the BA.4/5 spike protein harbors the F486V RBD mutation that can reduce monoclonal antibody sensitivity but had not yet been observed in other variants of interest or concern.^25^ Thus, newly emerged Omicron sublineages differ from BA.1 in key residues of the spike protein. Given their apparent growth advantages compared to BA.1 and the prevalent BA.2, the sublineages BA.2.12.1 and BA.4/5 are likely to be amongst the dominant variants of the SARS-CoV-2 pandemic in the near future.

### Reduced serum neutralizing activity against Omicron sublineages

To determine Omicron sublineage escape profiles, we measured neutralizing activity against the ancestral Wu01 strain as well as BA.1, BA.1.1, BA.2, BA.2.12.1, and BA.4/5 using lentivirus-based pseudovirus assays.^26,27^ In order to assess escape from polyclonal serum activity, we collected samples from two longitudinal cohorts of i.) SARS-CoV-2-convalescent individuals (*n*=20) and ii.) vaccinated health care workers (*n*=30) (**Extended Data Table 1**).^27,28^ In both cohorts, individuals received an mRNA vaccine booster immunization (BNT162b2) after a median of 14 months and 9 months following infection or two doses of BNT162b2, respectively.

Convalescent individuals had a median age of 51 years (interquartile range [IQR] 35-60) and were diagnosed with mild or asymptomatic SARS-CoV-2 infection. All individuals were infected during the initial phase of the pandemic between February and April 2020, prior to the emergence of designated viral variants of concern. Early post-infection samples (V1) were collected at a median of 48 days (IQR 34-58) after disease onset or diagnosis and neutralizing activity was assessed by determining the 50% inhibitory serum dilutions (ID_50_s) (**Fig. 2a**). Neutralization of the Wu01 strain was detected in all samples (100%) obtained early after infection, with individual ID_50_ values ranging from 16 to 2,607 (geometric mean ID_50_ [GeoMeanID_50_] of 264) (**Fig. 2b**). In contrast, serum activity against Omicron sublineages was strongly reduced and only detectable in 0-15% for BA.1 and BA1.1, and 45-50% for BA.2, BA.2.12.1, and BA.4/5 (**Fig. 2b**). However, analysis of sera obtained at a median of 33 days (IQR 27-54) after a single BNT162b2 booster immunization revealed neutralizing activity against all tested Omicron sublineages (**Fig. 2b, Extended Data Fig. 1a**) with GeoMeanID_50_s ranging from 1,456 against BA.4/5 to 2,103 against BA.2 (**Fig. 2b, Extended Data Fig. 1b,c**).

**Fig. 2.**
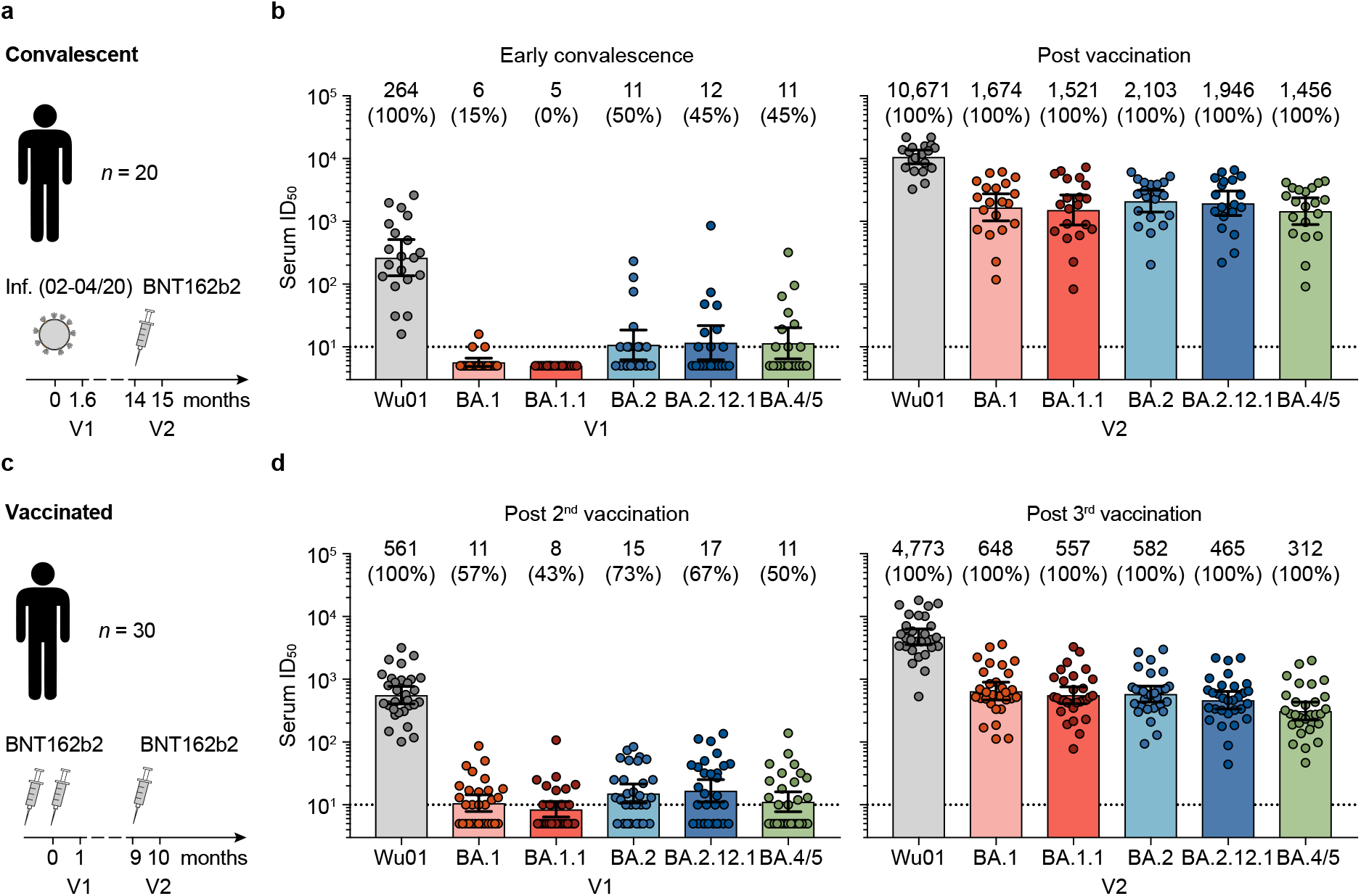
Omicron sublineage neutralizing serum activity in vaccinated and convalescent individuals. **a,** Study scheme in COVID-19-convalescent individuals. Samples were collected after infection occurring between February and April, 2020 (V1), and after a BNT162b2 booster immunization (V2). **b,** Fifty-percent inhibitory serum dilutions (ID_50_s) against Wu01, BA.1, BA.1.1, BA.2.12.1, and BA.4/5 determined by pseudovirus neutralization assays in convalescent individuals. Bars indicate geometric mean ID_50_s with 95% confidence intervals (CIs) at V1 (left) and V2 (right). Numbers indicate geometric mean ID_50_s and percentage of individuals with detectable neutralizing activity (ID_50_ >10) in parentheses. **c,** Study scheme in vaccinated individuals. Samples were collected after the second dose of BNT162b2 (V1) and after the third dose of BNT162b2 (V2). **d,** Serum ID_50_s against Wu01, BA.1, BA.1.1, BA.2.12.1, and BA.4/5 determined by pseudovirus neutralization assays in vaccinated individuals. Bars indicate geometric mean ID_50_s with 95% CIs at V1 (left) and V2 (right). Numbers indicate geometric mean ID_50_s and percentage of individuals with detectable neutralizing activity (ID_50_ >10) in parentheses. In **b** and **d**, ID_50_s below the lower limit of quantification (LLOQ, ID_50_ of 10; indicated by black dotted lines) were imputed to ½ LLOQ (ID_50_=5), and ID_50_s above the upper limit of quantification (21,870) were imputed to 21,871.

In addition, we determined Omicron sublineage neutralizing activity induced solely by vaccination (**Fig. 2c**). At a median of 28 days (IQR 27-32) after completion of the initial two-dose course of BNT162b2 (V1), Wu01 neutralizing serum activity was detected in all 30 individuals with a GeoMeanID_50_ of 561 (**Fig. 2d**). Although Omicron sublineage neutralization was detectable in 43-73% of vaccinated individuals at V1, GeoMeanID_50_s were very low (ranging from 8-17) (**Fig. 2d**). Follow-up samples obtained at a median of 29 days (IQR 26-35) after booster immunization showed 8-fold higher activity against Wu01 (GeoMeanID_50_ of 4,773) and strongly increased activity against all Omicron sublineages (27-to 70-fold GeoMeanID_50_ increases) (**Fig. 2d, Extended Data Fig. 1d-f**). Similar to the convalescent cohort, the lowest GeoMeanID_50_ after booster immunization was observed against the BA.4/5 sublineages (GeoMeanID_50_ of 312, **Fig. 2d**). Of note, amongst vaccinated individuals, the neutralizing activity against BA.4/5 was statistically significantly lower compared to that against BA.1, BA.1.1, and BA.2 (GeoMeanID_50_s of 648, 557, and 582, respectively) (**Extended Data Fig. 1e**).

We conclude that booster immunizations are critical to elicit neutralizing serum activity against all prevalent Omicron sublineages in vaccinated as well as convalescent individuals. Notably, BA.4/5 consistently demonstrated the lowest sensitivity of all sublineages against polyclonal serum. These results suggest increased antigenic escape of the recently emerged BA.4/5 sublineages mediated by the small number of spike protein differences relative to BA.2.

### Dissecting escape of Omicron sublineages

To decode Omicron escape from neutralizing antibodies on a molecular level, we produced and tested 158 monoclonal antibodies isolated from SARS-CoV-2-convalescent individuals against all sublineages. These included 67 randomly selected antibodies from the Coronavirus Antibody Database (CoV-AbDab),^29^ 79 antibodies isolated in our previous work,^30,31^ as well as 12 antibodies that were clinically tested. In total, this antibody panel originated from at least 43 different individuals out of 19 independent studies (**Fig. 3a**, **Extended Data Table 2**). All antibodies were identified in unvaccinated individuals that were infected in the early phase of the pandemic prior to the emergence of Omicron. The selection encompassed a broad spectrum of SARS-CoV-2 neutralizing antibodies with 92 V_H_/V_L_ combinations and a diverse range of CDR3 lengths and V gene mutations (**Fig. 3a**, **Extended Data Table 2**). Most antibodies targeted the RBD (97%) and the panel included previously described public clonotypes such as the V_H_3-53/3-66 subgroup (**Fig. 3a**, **Extended Data Table 2**).

**Fig. 3.**
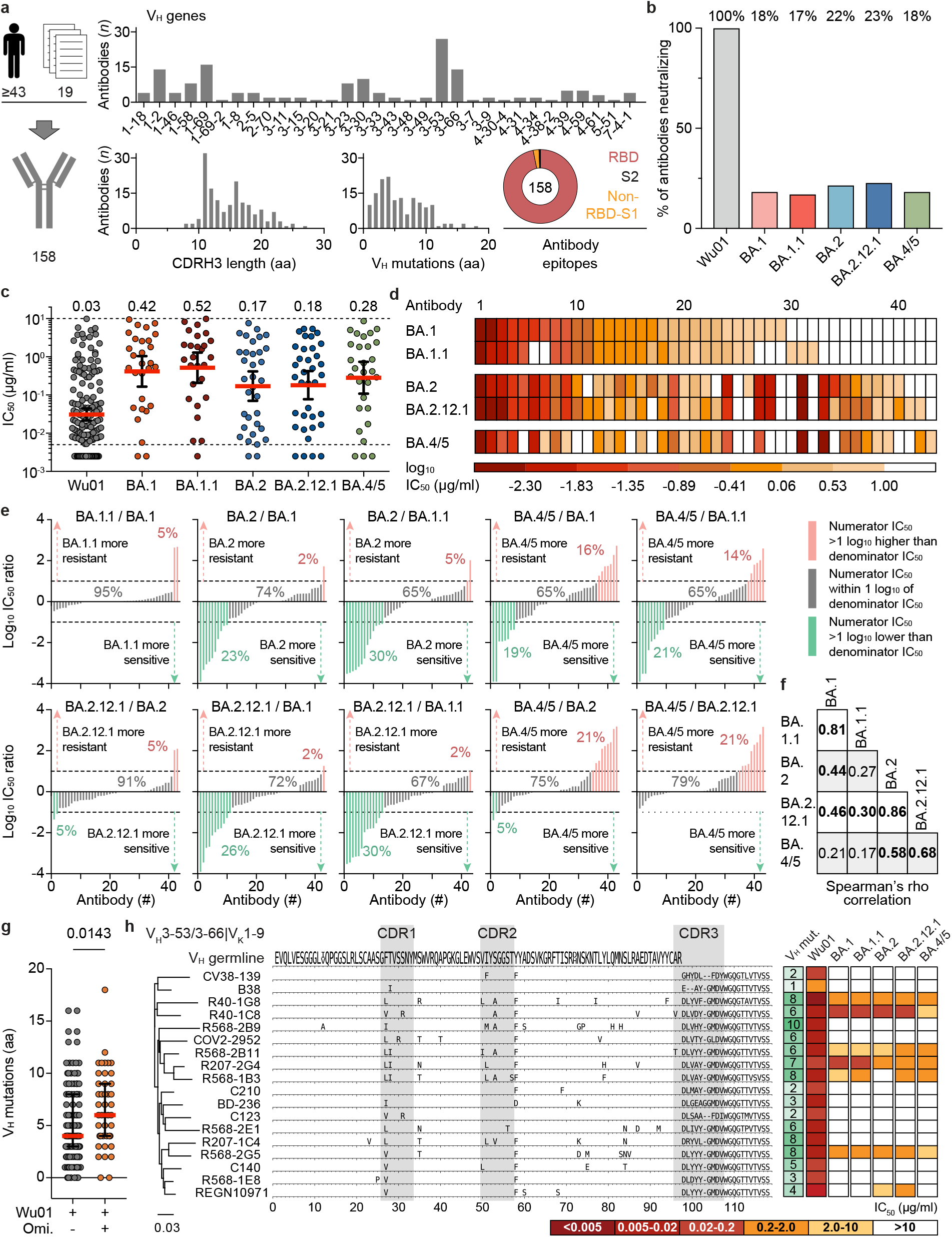
Determining Omicron sublineage immune escape using monoclonal antibodies. **a,** SARS-CoV-2 neutralizing monoclonal antibodies (*n*=158) derived from 19 studies and ≥43 convalescent individuals were analyzed. Bar charts indicate number of antibodies per heavy chain variable gene segment (V_H_), amino acid (aa) length of the heavy chain complementarity-determining region 3 (CDRH3), and number of V_H_ aa mutations relative to the V_H_ germline gene. Pie chart indicates antibody epitopes with slice sizes proportional to antibody fractions. RBD, receptor-binding domain. **b,** Fractions of antibodies from the panel (*n*=158) displaying activity (IC_50_ <10 µg/ml) against Wu01 and Omicron sublineages. **c,** IC_50_s of neutralizing antibodies against individual variants/sublineages (Wu01, *n*=158; BA.1, *n*=29; BA.1.1, *n*=27; BA.2, *n*=34; BA.2.12.1, *n*=*36*; BA.4/5, *n*=*29*). Solid lines depict geometric mean IC_50_s and 95% confidence intervals, and dashed lower and upper lines indicate lower (LLOQ, 0.005 µg/ml) and upper limit of quantification (ULOQ, 10 µg/ml), respectively. IC_50_s <LLOQ were imputed to ½ LLOQ (IC_50_=0.0025). **d,** Heat map depicting IC_50_s of the subset of antibodies (*n*=43) with neutralizing activity (IC_50_ <10 µg/ml) against at least one of the Omicron sublineages. Antibodies are sorted based on their potency against BA.1. **e,** Log_10_ IC_50_ ratios of antibodies neutralizing any Omicron sublineage for indicated sublineage comparisons. Within each panel, antibodies are sorted by increasing IC_50_ ratios. Ratios <-1 log_10_ are highlighted in green, ratios between -1 and 1 log_10_ in grey, and ratios >1 log_10_ in red. Numbers indicate relative fractions. IC_50_ <LLOQ were imputed to ½ LLOQ (IC_50_=0.0025) and IC_50_ values >ULOQ were imputed to 2x ULOQ (IC_50_=20). **f,** Spearman’s rank correlation coefficients (rho) for comparisons of antibodies with neutralizing capacity against any Omicron sublineage (*n*=43) as shown in **Fig. S2c**. Numbers in bold indicate values with *p*<0.05. **g,** Heavy chain V amino acid mutations of antibodies with neutralizing activity against Wu01 only and antibodies neutralizing both Wu01 and at least one omicron sublineage. Lines indicate medians and interquartile ranges. Groups were compared using the two-tailed Mann-Whitney U test. **h,** Phylogenetic tree and heavy chain sequence alignment of antibodies of the V_H_3-53/3-66|V_K_1-9 public clonotype. Letters indicate aa mutations relative to the V_H_ germline gene. Number of aa mutations from corresponding germline allele and neutralizing activity against Omicron sublineages are indicated on the right. Germline V_H_ represents the consensus of all identified germline alleles of depicted antibodies (V_H_3-53*01, V_H_3-53*04, and V_H_3-66*01). Number of aa mutations from corresponding germline allele and neutralizing activity against Wu01 and Omicron sublineages are indicated on the right.

While all 158 antibodies neutralized the Wu01 strain, only 18%, 17%, 22%, 23% and 18% remained active against BA.1, BA.1.1, BA.2, BA.2.12.1, and BA.4/5, respectively (**Fig. 3b, Extended Data Fig. 2a, Extended Data Table 2**). Moreover, amongst antibodies with retained neutralizing activity, the overall potency against Omicron sublineages was 6-to 14-fold lower compared to Wu01 with GeoMeanIC_50_s of 0.42 (BA.1), 0.52 (BA.1.1), 0.17 (BA.2), 0.18 (BA.2.12.1), and 0.28 µg/ml (BA.4/5) (**Fig. 3c**).

The neutralization profiles revealed both commonalities and diversity in antibody sensitivity of the different Omicron sublineages (**Fig. 3d-f, Extended Data Fig. 2b**). Between the most closely related sublineages (BA.1 and BA.1.1; BA.2 and BA.2.12.1), only small differences in antibody activity were observed (r_s_ of 0.81 and 0.86, respectively) (**Fig. 3e,f** **Extended Data Fig. 2c**). In contrast, comparisons between the primary sublineages BA.1, BA.2 and BA.4/5 demonstrated higher degrees of variation in antibody susceptibility (**Fig. 3e,f**). For example, BA.2 was more resistant (>1 log_10_ IC_50_ difference) than BA.1 to only a single antibody (2%), but more sensitive to ten antibodies (23%) (**Fig. 3e**). Moreover, higher fractions of antibodies showed >10-fold higher resistance against BA.4/5 relative to other variants (**Fig. 3e**). While the comparison of BA.4/5 to BA.1 and BA.1.1 revealed heterogeneity in sensitivity to the antibody panel with no correlation, the sensitivity of BA.4/5 and BA.2 or BA.2.12.1 more strongly correlated (r_s_ of 0.58 and 0.68, respectively). Importantly, however, whereas BA.4/5 were more sensitive to only 0-5% of antibodies compared to BA.2 and BA.2.12.1, BA.4/5 showed higher resistance to 21% of the antibodies (**Fig. 3e**).

Based on the analysis of the sublineage neutralization profiles, three prevalent classes of Omicron neutralizing antibodies became apparent: i.) antibodies with broadly comparable activity against Omicron sublineages (21/43, 49%), ii.) antibodies with strongly reduced activity against BA.1 and BA.1.1 compared to BA.2, BA.2.12.1, and BA.4/5 (7/43, 16%), and iii.) antibodies with strongly reduced activity against BA.4/5 compared to BA.1, BA.1.1, as well as BA.2 and/or BA.2.12.1 (6/43, 14%).

We conclude that in contrast to the limited differences in sensitivity to polyclonal serum, Omicron sublineages considerably differ in their sensitivity to monoclonal antibodies. Importantly, BA.4/5 demonstrated a strong bias to higher resistance compared to the prevalent BA.2 sublineages.

### Minor sequence variations critically affect antibody activity

Antibodies with Omicron neutralizing activity carried a modestly higher number of amino acid mutations in the heavy and light chain variable gene compared to antibodies only neutralizing Wu01 (median of 6 vs. 4 and 4 vs. 3 mutations; *p*=0.0143 and *p*=0.0170, respectively), suggesting that higher sequence diversification might be overall favorable for Omicron neutralization (**Fig. 3f, Extended Data Fig. 2e**).

Antibody responses against SARS-CoV-2 are highly convergent across individuals based on the identification of public clonotypes with conserved sequence characteristics and neutralization mechanisms.^32,33,34^ Amongst the analyzed antibody panel, 18 sequences from 11 individuals could be assigned to the prominent V_H_3-53/3-66|V_K_1-9 clonotype (**Fig. 3g, Extended Data Fig. 2f**).^30,35,36^ Although these antibodies are highly conserved on a sequence level, they substantially differed in their Omicron neutralizing capacity (**Fig. 3g, Extended Data Fig. 2f**). For example, antibodies R207-1C4 and R568-2G5 harbor eight amino acid mutations in their V_H_ gene segment (without CDRH3) of which five are at the same position and three are identical. While both antibodies had similar Wu01 neutralizing activity, R207-1C4 did not neutralize any Omicron sublineage, whereas R568-2G5 showed neutralizing activity against all variants. Notably, antibody C140, another member of this clonotype which has the identical CDRH3 motif as R568-2G5, did not neutralize any Omicron variant.

We conclude that minimal variation in antibody sequences can tip the scale between Omicron neutralization and resistance. Thus, sequence- or antibody class-based predictions of Omicron neutralization are difficult and experimental assessment of individual monoclonal antibody activity remains essential.

### Impact on clinical and novel monoclonal antibodies

SARS-CoV-2 neutralizing monoclonal antibodies can reduce morbidity and mortality in infected individuals and are critical for passive immunization to protect individuals that do not mount an adequate immune response upon vaccination.^2,3,37–41^ To determine how spike protein mutations of Omicron sublineages affect antibodies in clinical use, we analyzed 9 monoclonal antibodies that received authorization for clinical use (**Fig. 4a,b**) and 9 that are advanced in clinical development (**Fig. 4b, Extended Data Fig. 3**). All antibodies target the RBD of the SARS-CoV-2 spike protein and were tested against Wu01 and the Omicron sublineages (**Fig. 4b**).

**Fig. 4.**
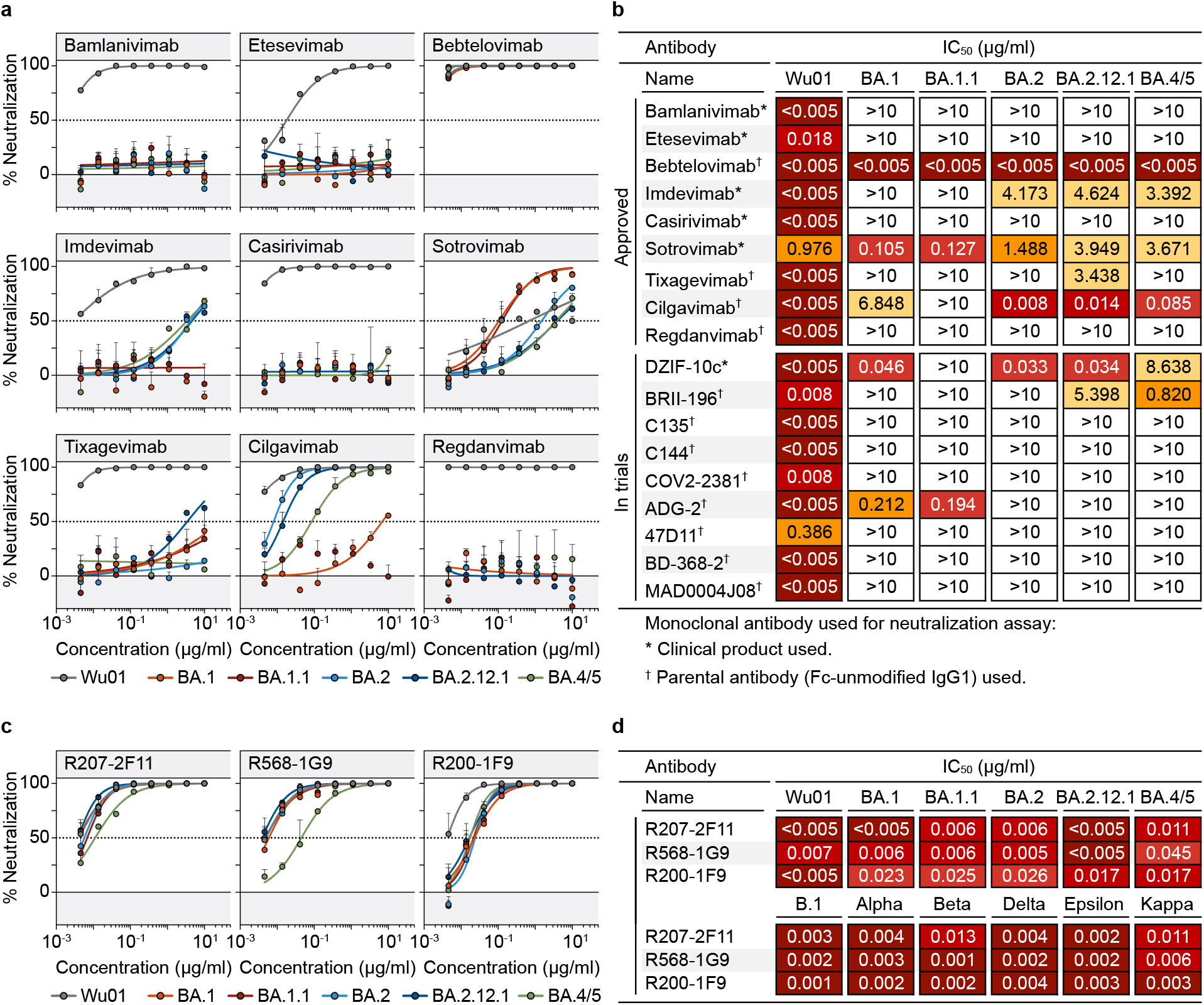
Omicron sublineage neutralizing activity of monoclonal antibodies in clinical use. **a,** Dose response curves showing percentage neutralization of monoclonal antibodies against Wu01 and Omicron sublineages in pseudovirus neutralization assay. Circles show averages and error bars indicate standard deviation. Dotted lines indicate 50% neutralization (IC_50_). **b,** IC_50_s of monoclonal antibodies with current or previous authorization for clinical use or in clinical development. Symbols indicate whether clinical products or parental antibodies produced as human IgG1 were used. **c,** Dose response curves showing percentage neutralization of novel monoclonal antibodies against Wu01 and Omicron sublineages in pseudovirus neutralization assay. Circles show averages and error bars indicate standard deviation. Dotted lines indicate 50% neutralization (IC_50_). **d,** IC_50_s of novel monoclonal antibodies shown in **c** against Wu01 and Omicron sublineages (upper rows) and previously circulating variants (lower rows, as previously determined_30_).

Most antibodies showed highly potent neutralizing activity against Wu01 with IC_50_s below 0.005 µg/ml (**Fig. 4b**). Less potent and incomplete Wu01 neutralizing activity was observed for sotrovimab, consistent with previous reports showing reduced activity against pseudoviruses lacking the dominant D614G spike mutation (**Fig. 4a**).^42,43^ Only five (28%) out of the 18 tested antibodies neutralized BA.1 with IC_50_s <10 µg/ml, and of these, three had strongly reduced potency (**Fig. 4b**). While the antibody neutralization profile of BA.1.1 was generally similar to that of BA.1, there were some differences. For example, DZIF-10c neutralized BA.1 with an IC_50_ of 0.046 µg/ml but completely lost activity against BA.1.1 (**Fig. 4b, Extended Data Fig. 3**).

Although the number of antibodies in clinical use with neutralizing activity against BA.2 (5 out 18, 28%), BA.2.12.1 (7 out 18, 39%), and BA.4/5 (6 out 18, 33%) was similarly small, the neutralization profiles did differ (**Fig. 4b**). For example, cilgavimab neutralized BA.2 and BA.2.12.1 with high potency (IC_50_s of 0.008 and 0.014 µg/ml, respectively) but was >480-fold less potent against BA.1 and BA.1.1 (**Fig. 4a,b**). While, cilgavimab maintained high potency against BA.4/5 (IC_50_ of 0.085 µg/ml), the activity of antibody DZIF-10c against BA.4/5 was strongly reduced when compared to BA.2 and BA.2.12.1 (8.64 vs. 0.03 µg/ml) (**Fig. 4b**). In contrast, whereas antibody imdevimab showed no appreciable activity against BA.1 and BA.1.1, neutralization of BA.2, BA.2.12.1, and BA.4/5 was detectable at low levels (**Fig. 4a,b**).

Out of all clinical antibodies tested, only bebtelovimab maintained high neutralization potency against all Omicron sublineages (IC_50_ <0.005 µg/ml; **Fig. 4a,b**). To identify additional promising candidates, we examined antibodies previously isolated from individuals with ‘elite’ IgG neutralizing activity.^30^ Remarkably, antibodies R207-2F11, R568-1G9, and R200-1F9 demonstrated highly potent neutralizing activity against previously circulating variants (Wu01, B.1, Alpha, Beta, Delta, Epsilon, and Kappa), as well as against BA.1, BA.1.1, BA.2, BA.2.12.1, and BA.4/5 with IC_50_s <0.05 µg/ml (**Fig. 4d, Extended Data Table 2**). These antibodies were identified from three different convalescent individuals within weeks of infection during the early phase of the pandemic.^30^ Thus, the results highlight the capability of the immune system to generate potent antibodies against the highly divergent Omicron variants in response to contact with the ancestral spike protein.

We conclude that the prevalent and newly emerged Omicron sublineages, including BA.2.12.1 and BA.4/5, escape from most monoclonal antibodies in clinical use. Omicron sensitivity to these antibodies can, however, strongly differ on the sublineage level.

## Discussion

Antibody-mediated immunity is a critical component of prophylactic and therapeutic measures against SARS-CoV-2 infection.^2,3,44,45^ Throughout the COVID-19 pandemic, it has however been confronted by the periodic emergence of antibody escape variants developing in the context of increasing population immunity.^46^ Omicron sublineages with diverse spike proteins gaining prevalence suggest further increase in immune escape and/or transmissibility.^20,21,47^ Thus, establishing the impact of the recent BA.2.12.1 and BA.4/5 sublineages on polyclonal and monoclonal immunity is critical to guide antibody-mediated strategies for prevention and treatment.

Our results demonstrate that Wu01-based mRNA vaccine boosters are effective in eliciting serum neutralizing activity against the diverse Omicron sublineages. However, this activity is strongly limited compared to Wu01 neutralization and lower BA.4/5 neutralizing titers indicate even more pronounced immune evasion relative to previous Omicron sublineages. Interestingly, the relative reduction in activity against BA.4/5 after booster immunization was higher in infection-naïve than in convalescent individuals. This might indicate differences in the breadth and potency of the neutralizing antibody response elicited by infection only compared to those induced by “hybrid immunity”.

While the differences in polyclonal serum sensitivity between the Omicron sublineages were overall modest, our analysis of an extended antibody panel revealed distinct patterns in sublineage sensitivity on a monoclonal level. Comparing Omicron sublineages revealed the largest antigenic distance between BA.4/5 and BA.1/BA.1.1 as indicated by the lack of correlation in antibody sensitivity and notable fractions of antibodies showing higher sensitivity to one or the other sublineage. Whereas BA.4/5 and BA.2/BA.2.12.1 showed more similar antibody susceptibility profiles, almost all differences manifested as higher antibody resistance of BA.4/5.

Notably, minor differences in antibody sequence strongly affected Omicron activity of antibodies with high potency against Wu01, highlighting the relative difficulty of inducing antibodies with optimized Omicron neutralization after single contact with Wu01-related spike. Accordingly and consistent with previous analyses on BA.1 and BA.2, ^48,49^ almost all antibodies in clinical investigation showed strongly reduced or abolished activity against all Omicron sublineages. However, we identify novel antibodies isolated after ancestral infection with remarkable activity against all tested variants that may provide options for immune-mediated treatment and prevention in the Omicron era of the pandemic.

Our results suggest that outcompetition of BA.1(.1) by BA.2 may have been driven mostly by higher transmissibility rather than by immune evasion. In contrast, higher antigenic distance and antibody resistance of BA.4/5 may support its expansion after the initial Omicron wave. The rapid emergence of Omicron sublineages with reduced antibody susceptibility highlights the challenges posed by constant viral evolution and the critical need for genomic surveillance and sensitivity assessments to inform on antibody-based prophylactic and therapeutic measures.

## Acknowledgments

We are grateful to all study participants for their dedication to our research. We thank F. Dewald, L. Gieselmann, B. Kurt, C. Lehmann, P. Mayer, N. Riet, S. Salomon, M. Schlotz, R. Schröder, R. Stumpf, and H. Wüstenberg, as well as the members of the EICOV/COVIM Study Group (Y. Ahlgrimm, L. Bardtke, K. Behn, N. Bethke, H. Bias, D. Briesemeister, C. Conrad, V. M. Corman, C. Dang-Heine, S. Dieckmann, C. Eroglu, D. Frey, J.-A. Gabelich, J. Gerdes, U. Gläser, A. Hetey, L. Hasler, E. T. Helbig, D. Hillus, W. Hirst, A. Horn, C. Hülso, S. Jentzsch, C. von Kalle, L. Kegel, A. Krannich, W. Koch, P. Kopankiewicz, P. Kroneberg, H. Le, L. J. Lippert, C. Lüttke, P. de Macedo Gomes, B. Maeß, C. Matthaiou, J. Michel, A. Nitsche, A.-M. Ollech, C. Peiser, A. Pioch, C. Pley, K. Pohl, C. Rubisch, L. Ruby, A. Sanchez Rezza, I. Schellenberger, V. Schenkel, J. Schlesinger, S. Schmidt, G. Schwanitz, T. Schwarz, J. Seybold, A. Solarek, A. Stege, S. Steinbrecher, P. Stubbemann, C. Thibeault, and D. Treue) for sample acquisition and processing; L. Buchholz and M. Wunsch for help with antibody production and preparation; and J.J. Malin for providing clinical antibody stocks. We thank all laboratories obtaining and submitting SARS-CoV-2 sequencing information to GISAID for facilitating rapid analyses of global variant distributions. This work was supported by grants from COVIM: NaFoUniMedCovid19 (FKZ: 01KX2021) (to L.E.S. and F. Klein), the Federal Institute for Drugs and Medical Devices (V-2021.3 / 1503_68403 / 2021–2022) (to L.E.S. and F. Kurth), the German Center for Infection Research (DZIF) (to F. Klein), and the Deutsche Forschungsgemeinschaft (DFG) CRC1310 (to C.K. and F. Klein).

## Author contributions

Conceptualization, H.G., K.V., C.K., and F. Klein; methodology, H.G., K.V., L.E.S., F. Kurth, C.K., and F. Klein; investigation, H.G. and K.V.; resources, P.T-L., M.K., M.Z., F.M., H.J., M.A., and P.S.; formal analysis, H.G., K.V., and C.K.; writing-original draft, H.G., K.V., C.K., and F. Klein; writing-review and editing, M.K., P.T-L., and F. Kurth; visualization, H.G., K.V., C.K., and F. Klein; supervision, L.E.S., F. Kurth, and F. Klein; funding acquisition, L.E.S., F. Kurth, and F. Klein.

## Extended Data Figure legends

**Extended Data Fig. 1. Serum neutralization of Omicron sublineages.**

**a,** Serum ID_50_s against Wu01 and Omicron sublinages in the cohort of convalescent individuals after infection (V1) and BNT162b2 booster immunization (V2) as in **Fig. 2**. Lines connect ID_50_s of individual participants at V1 and V2. Dashed lines indicate lower limit of quantification (LLOQ, ID_50_=10). **b,** Numbers in black indicate Spearman’s rank correlation coefficients (rho) for comparisons at V2 shown in **c**. Numbers in red indicate *p* values determined by Friedman test followed by Dunn’s multiple comparison test for comparisons at V2, with statistically significant differences indicated in bold. **c,** Correlation plots of log_10_ serum ID_50_s against indicated viruses in convalescent individuals at V2. **d,** Serum ID_50_s against Wu01 and Omicron sublineages in the cohort of BNT162b2-vaccinated individuals after the second (V1) and the third vaccine dose (V2) as in **Fig. 2**. Lines connect ID_50_s of individual participants at V1 and V2. Dashed lines indicate lower limit of quantification (LLOQ, ID_50_=10). **e,** Numbers in black indicate Spearman’s rank correlation coefficients (rho) for comparisons at V2 shown in **f**. Numbers in red indicate *p* values determined by Friedman test followed by Dunn’s multiple comparison test for comparisons at V2, with statistically significant differences indicated in bold. **f,** Correlation plots of log_10_ serum ID_50_s against indicated viruses in vaccinated individuals at V2. In **a** and **d**, Serum ID_50_s <LLOQ were imputed to ½ LLOQ (ID_50_=5).

**Extended Data Fig. 2. Neutralization profile of Omicron neutralizing antibodies.**

**a,** Spider plot of IC_50_s for all antibodies against Wu01 and Omicron sublineages. Antibodies are sorted arbitrarily but equally for each virus. Circles indicate IC_50_s (from outer to inner: 0.005, 0.05, 0.5, and 5 µg/ml). **b,** Bar charts of antibodies with neutralizing activity (IC_50_ <10 µg/ml) against any Omicron sublineage (*n*=43). In each chart, antibodies are sorted by BA.1-neutralizing activity. Dotted lines indicate lower (LLOQ, 0.005 µg/ml) and upper limits of quantification (ULOQ; 10 µg/ml). **c,** IC_50_ correlation plots for antibodies with neutralizing capacity against any Omicron sublineage (*n*=43). Colors indicate epitopes as in **Fig. 3a**. Grey dashed lines represent identity lines and black dashed lines indicate limits of quantification. In **a-c**, IC_50_s <LLOQ were imputed to ½ LLOQ (IC_50_=0.0025) and IC_50_s >ULOQ were imputed to 2x ULOQ (IC_50_=20). **d,** Light chain V amino acid mutations of antibodies with neutralizing activity against Wu01 only and antibodies neutralizing both Wu01 and at least one omicron sublineage. Lines indicate medians and interquartile ranges and groups were compared using the two-tailed Mann-Whitney U test. **e,** Light chain sequence alignment of antibodies of the V_H_3-53/3-66|V_K_1-9 public clonotype. Letters indicate amino acid (aa) mutations relative to the light chain germline gene. Germline V_K_ represents the consensus of all identified germline alleles of the depicted antibodies. Number of aa mutations from corresponding germline allele and neutralizing activity against Wu01 and Omicron sublineages are indicated on the right.

**Extended Data Fig. 3. Omicron sublineage neutralizing activity of monoclonal antibodies in clinical testing.**

Dose response curves showing % neutralization of monoclonal antibodies against Wu01 and Omicron sublineages in a pseudovirus neutralization assay. Circles show averages and error bars indicate standard deviation. Dotted lines indicate 50% neutralization (IC_50_).

## Extended Data Tables

Extended Data Table 1: Study cohorts

Extended Data Table 2: Monoclonal antibody panel analyses

## Methods

### Study cohort and sample collection

Serum samples from COVID-19-convalescent individuals were collected at the University Hospital Cologne under study protocols approved by the ethics committee (EC) of the Medical Faculty of the University of Cologne (16-054 and 20-1187). Between April and May, 2020, individuals with a history of SARS-CoV-2 infection confirmed by polymerase chain reaction (documented through a written test certificate or as reported to study investigators by the participants) were enrolled within eight weeks of symptom onset and/or diagnosis. As all participants were enrolled early during the pandemic (i.e., prior to the emergence of variants of concern, as designated by the World Health Organization), most individuals are likely to have been infected with an early viral strain similar to Wu01. Participants were followed longitudinally to analyze long-term immunity to SARS-CoV-2. Follow-up samples after booster immunization were collected between May and August, 2021. No additional SARS-CoV-2 infections between the sampling time points were reported by any of the participants.

Serum samples from vaccinated individuals were collected under protocols approved by the EC of Charité - Universitätsmedizin Berlin (EICOV, EA4/245/20) as well as the EC of the Federal State of Berlin and the Paul Ehrlich Institute (COVIM, EudraCT-No. 2021-001512-28). Study participation irrespective of medical conditions was offered to health-care workers vaccinated at the Charité – Universitätsmedizin (Berlin, Germany). All serum samples were tested for antibodies targeting the SARS-CoV-2 nucleocapsid using the SeraSpot Anti-SARS-CoV-2 IgG microarray-based immunoassay (Seramun Diagnostica). Samples from individuals with a history of SARS-CoV-2 infection, a positive SARS-CoV-2 nucleic acid amplification test (performed at sampling), or detectable anti-nucleocapsid antibodies were not included in this analysis. Serum samples after two vaccinations were collected in February and March 2021 (for 29/30 participants; from one participant, the early serum sample was obtained in July 2021); samples after booster immunization were collected in December 2021 and January 2022.

All study participants provided written informed consent. Vaccinations in both cohorts were performed as part of routine care outside of the observational studies. Selection of participants and samples for analysis was based on receipt of identical vaccines and comparable sampling time points relative to vaccinations. Serum samples were collected after centrifugation and stored at -80°C until analysis.

### SARS-CoV-2 pseudovirus constructs

All SARS-CoV-2 spike proteins were expressed using codon-optimized expression plasmids. Wu01 (EPI_ISL_406716), BA.2.12.1, and BA.4/5 pseudoviruses were produced using an expression plasmid that incorporated a C-terminal deletion of 21 cytoplasmic amino acids that result in increased pseudovirus titers. Expression plasmids for Omicron sublineage spike proteins were produced by assembling and cloning codon-optimized overlapping gene fragments (Thermo Fisher) into the pCDNA3.1/V5-HisTOPO vector (Thermo Fisher) using the NEBuilder HiFi DNA Assembly Kit (New England Biolabs), and included the full spike protein amino acid sequences with the following amino acid changes relative to Wu01:

BA.1: A67V, Δ69-70, T95I, G142D, Δ143-145, N211I, Δ212, ins215EPE, G339D, S371L, S373P, S375F, K417N, N440K, G446S, S477N, T478K, E484A, Q493R, G496S, Q498R, N501Y, Y505H, T547K, D614G, H655Y, N679K, P681H, N764K, D796Y, N856K, Q954H, N969K, and L981F.

BA.1.1: As for BA.1 with an additional R346K mutation.

BA.2: T19I, Δ24-26, A27S, A67V, G142D, V213G, G339D, S371F, S373P, S375F, T376A, D405N, R408S, K417N, N440K, S477N, T478K, E484A, Q493R, Q498R, N501Y, Y505H, D614G, H655Y, N679K, P681H, N764K, D796Y, Q954H, N969K.

BA.2.12.1: As for BA.2 with additional L452Q and S704L mutations.

BA.4/5: As for BA.2 with additional Δ69-70, L452R, and F486V mutations, but lacking the Q493R mutation.

All plasmid sequences were verified by sequencing.

### Monoclonal antibodies

Monoclonal antibodies previously isolated in our lab had been obtained by single cell-sorting of SARS-CoV-2 spike-specific B cells followed by reverse transcription, PCR amplification and cloning of antibody variable regions.^30,31^ For monoclonal antibodies derived from the CoV-AbDab ^29^, variable region amino acid sequences were reverse translated with the reverse translate tool from the Sequence Manipulation Suite^50^ using the *Homo sapiens* codon table obtained from the Codon Usage Database,^51^ and sequences were ordered as gene fragments from Integrated DNA Technologies (IDT) with 5’ and 3’ overhangs. The variable regions were inserted into heavy and light chain expression plasmids^52^ by sequence- and ligation-independent cloning (SLIC). For antibodies ADG-2, COV2-2130 (cilgavimab), COV2-2196 (tixagevimab), COV2-2381, MAD0004J08, and P2C-1F11 (BRII-196), gene fragments based on the nucleotide sequences published in GenBank were ordered at IDT and cloned as above. For antibodies C135, CT-P59 (regdanvimab), and LY-CoV1404 (bebtelovimab), gene fragments based on antibody structures deposited in the Protein Data Bank (accession nos. 7K8Z, 7CM4, and 7MMO) were ordered at IDT after codon optimization using the IDT Codon Optimization Tool and cloned as above. For antibodies 47D11, BD-368-2, C144, and P2B-2F6, amino acid sequences were derived from CoV-AbDaB, corresponding nucleotide sequences generated and codon-optimized using the IDT Codon Optimization Tool, and gene fragments cloned as above.

Monoclonal antibody production was performed using 293-6E cells (National Research Council of Canada) by co-transfection of heavy and light chain expression plasmids using 25 kDa branched polyethylenimine (Sigma-Aldrich). Culture supernatants were harvested after an incubation period of 6-7 days at 37°C and 6% CO2 under constant shaking in FreeStyle Expression Medium supplemented with penicillin (20 U/ml) and streptomycin (20 µg/ml) (all Thermo Fisher). Clarified cell supernatants were incubated with Protein G Sepharose 4 FastFlow (Cytiva) overnight at 4°C . After centrifugation, antibodies bound to Protein G beads were eluted in chromatography columns (Bio-Rad) using 0.1 M glycine (pH=3.0) and buffered in 1 M Tris (pH=8.0). Buffer exchange to PBS was performed using centrifugal filter units (Millipore). For antibodies bamlanivimab, casirivimab, DZIF-10c, etesevimab, imdevimab, and sotrovimab, aliquots from clinical stocks were used.

### Pseudovirus neutralization assays

Neutralization assays were performed using lentivirus-based pseudoviruses and ACE2-expressing 293T cells.^26,27^ Pseudovirus particle production was performed in HEK293T cells by co-transfection of individual expression plasmids encoding for the SARS-CoV-2 spike protein, HIV-1 Tat, HIV-1 Gag/Pol, HIV-1 Rev, and luciferase-IRES-ZsGreen using FuGENE 6 Transfection Reagent (Promega). Culture supernatants were exchanged with fresh medium (high glucose DMEM supplemented with 2 mM L-glutamine, 100 IU/ml penicillin, 100 µg/ml streptomycin, 1 mM sodium pyruvate (all Thermo Fisher), and 10% FBS (Sigma-Aldrich)) 24 h post transfection. Pseudovirus-containing supernatants were harvested between 48-72 h after transfection, centrifuged, clarified using a 0.45 µm filter, and stored at -80°C. Pseudoviruses were titrated by infection of 293T-ACE2 cells and luciferase activity was determined after a 48-hour incubation at 37°C and 5% CO_2_ by addition of luciferin/lysis buffer (10 mM MgCl_2_, 0.3 mM ATP, 0.5 mM coenzyme A, 17 mM IGEPAL CA-630 (all Sigma-Aldrich), and 1 mM D-Luciferin (GoldBio) in Tris-HCL) using a microplate reader (Berthold).

Serum samples were heat-inactivated at 56°C for 45 min before use. Three-fold serial dilutions of serum (starting at 1:10) and monoclonal antibodies (starting at 10 µg/ml) were prepared in culture medium and co-incubated with pseudovirus supernatants for one hour at 37°C and 5% CO_2_ prior to addition of 293T-ACE2 cells. Following a 48-hour incubation at 37°C and 5% CO_2_, luciferase activity was determined as described above. Average background relative light units (RLUs) of non-infected cells were subtracted, and serum ID_50_s and antibody IC_50_s were determined as the serum dilutions and antibody concentrations resulting in a 50% RLU reduction compared to the average of virus-infected untreated controls cells using a non-linear fit model plotting an agonist vs. normalized dose response curve with variable slope using the least squares fitting method in Prism 7.0 (GraphPad). All serum and monoclonal antibody samples were tested in duplicates. Imputation rules for values outside the limits of quantification are described below (see Statistical methods).

### SARS-CoV-2 neutralizing antibody panel and sequence analysis

The panel of 158 SARS-CoV-2 neutralizing monoclonal antibodies isolated from SARS-CoV-2 convalescent individuals included in the analysis in **Fig. 3** is based on 79 antibodies obtained in our previous work,^30,31^ 67 randomly selected (retrieved on January 1, 2021) human SARS-CoV-2 neutralizing antibodies deposited at CoV-AbDab,^29^ and 12 antibodies in clinical use or development. We did not include five of the antibodies in clinical development shown in **Fig. 4** into this analysis, as they were obtained from individuals infected with SARS-CoV (ADG-2, sotrovimab), from immunized mice harboring human immunglobulin gene repertoires (47D11, casirivimab), or using phage display technology that does not ensure native pairing of antibody heavy and light chains (regdanvimab).

Antibody amino acid sequences were annotated with IgBLASTp^53^ based on the IMGT database.^54^ For sequence statistics, top V gene calls were counted without individual alleles, CDR3 lengths are reported according to the IMGT numbering system and numbers of V mutations refer to the top V gene call from IgBLASTp. Phylogenetic analysis of antibodies belonging to the V_H_3-53/3-66|V_K_1-9 public clonotype was performed by alignment of amino acid sequences with the MAFFT algorithm^55^ via the EMBL-EBI search and sequence analysis tools API^56^ and the Tree Builder tool from Geneious Prime 2020.0.4 (Biomatters) using the Jukes-Cantor distance model for tree building with the neighbour-joining method without resampling. Data aggregation and visualization was performed with the Python libraries pandas (v1.1.5), NumPy (v1.19.2), SciPy (v1.5.2), Matplotlib (v3.3.4) with Python (v3.6.8), as well as Microsoft Excel 2011 for Mac (v14.7.3), and Adobe Illustrator.

### Visualization of SARS-CoV-2 spike amino acid changes

Amino acid changes relative to the Wu01 spike protein were visualized on a cryo-electron microscopy 3D-reconstruction of the SARS-CoV-2 spike protein (PDB ID: 6XR8, ref. ^57^) using ChimeraX (v. 1.3).^58,59^

### SARS-CoV-2 variant distribution

Numbers of weekly cases as curated by the COVID-19 Data Repository by the Center for Systems Science and Engineering (CSSE) at Johns Hopkins University^60^ was retrieved from http://ourworldindata.org (accessed on May 24, 2022). GISAID-curated clade and lineage statistics of sequences submitted to the GISIAD database^61–63^ were retrieved from GISAID (accessed on May 24, 2022) and frequency of individual variants was plotted as fraction of all submitted sequences per week and variant. For visualization, all BA.1 sublineages except for BA.1.1 (and sublineages of BA.1.1) were classified as BA.1, and all BA.2 sublineages except for BA.2.12.1 were classified as BA.2

## Statistical Methods

For graphical representation and statistical evaluation of serum samples in **Fig. 2** and **Extended Data Fig. 1**, samples that did not achieve 50% inhibition at the lowest tested dilution of 10 (lower limit of quantification, LLOQ) were imputed to ½ of the LLOQ (ID_50_=5) and serum samples with ID_50_ >21,870 (upper limit of quantification) were imputed to ID_50_=21,871. For graphical representation and statistical analysis of monoclonal neutralizing antibodies in **Fig. 3** and **Extended Data Fig. 2**, IC_50_ values of antibodies with an IC_50_ <0.005 µg/ml (LLOQ) were imputed to ½ LLOQ (IC_50_=0.0025), and IC_50_ values >10 µg/ml (ULOQ) were imputed to 2x ULOQ (IC_50_=20 µg/ml).

Testing for statistical significance of differences in serum neutralization titers against different variants/sublinages was performed with the Friedman test with Dunn’s multiple comparison using Prism 7.0 (GraphPad). Spearman’s rank correlation coefficients (Rho) were determined using Prism 7.0 (GraphPad). Amino acid mutations relative to germline between antibodies neutralizing Wu01 only and those neutralizing any Omicron sublineage were compared with two-sided Mann-Whitney U tests using Prism 7.0 (GraphPad).

## Data and code availability

Requests for data or materials should be directed to the corresponding author and may be subject to restrictions based on data and privacy protection regulations and/or may require a Material Transfer Agreement (MTA). The paper does not report original code.

