## Supplemental Information for "Delineating antibody escape from Omicron sublineages"

### Extended Data Figure 1

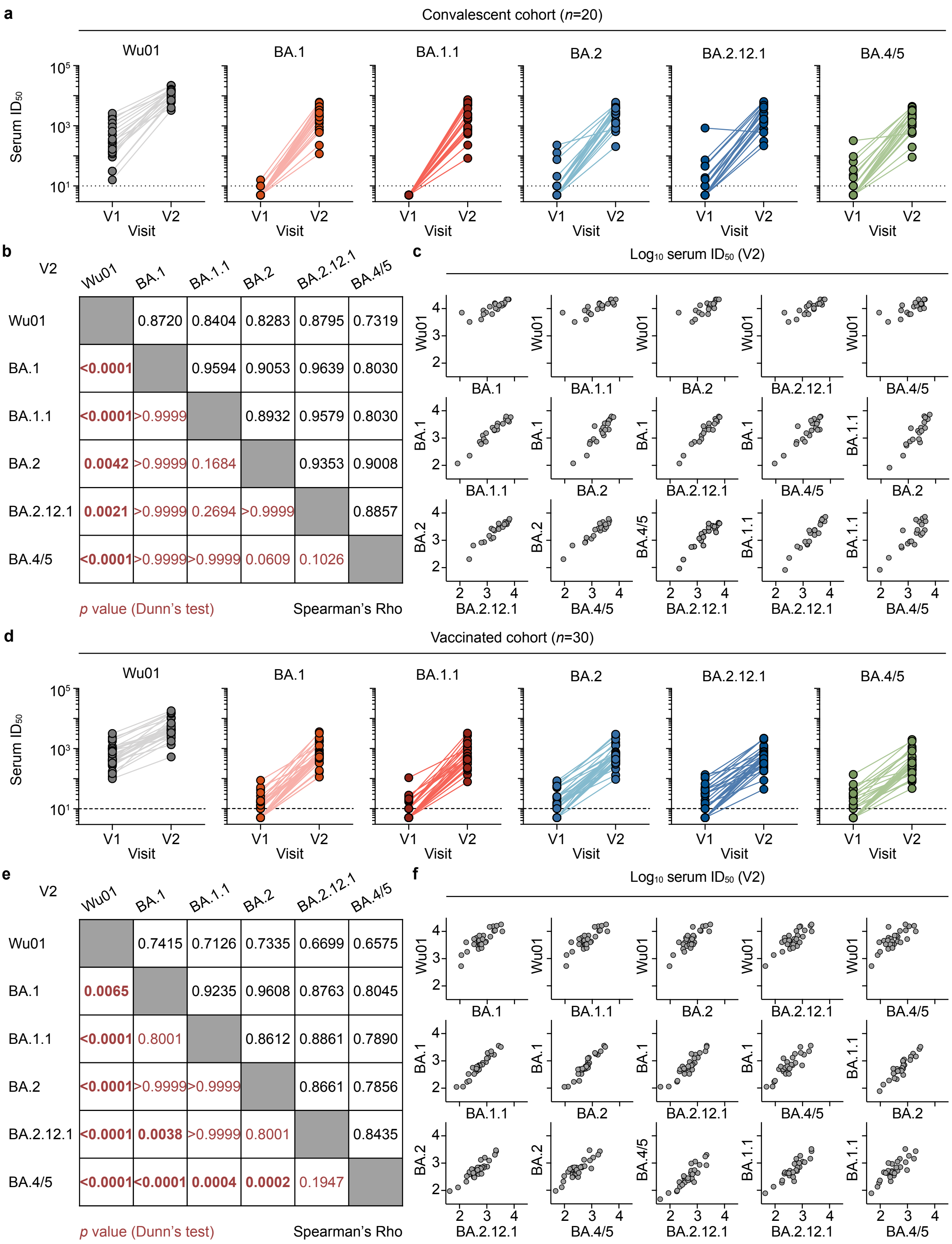


**b**

| V2 | Wu01 | BA.1 | BA.1.1 | BA.2 | BA.2.12.1 | BA.4/5 |
|---|---|---|---|---|---|---|
| Wu01 | | 0.8720 | 0.8404 | 0.8283 | 0.8795 | 0.7319 |
| BA.1 | **<0.0001** | | 0.9594 | 0.9053 | 0.9639 | 0.8030 |
| BA.1.1 | **<0.0001** | >0.9999 | | 0.8932 | 0.9579 | 0.8030 |
| BA.2 | **0.0042** | >0.9999 | 0.1684 | | 0.9353 | 0.9008 |
| BA.2.12.1 | **0.0021** | >0.9999 | 0.2694 | >0.9999 | | 0.8857 |
| BA.4/5 | **<0.0001** | >0.9999 | >0.9999 | 0.0609 | 0.1026 | |

*p* value (Dunn's test) — Spearman's Rho

**e**

| V2 | Wu01 | BA.1 | BA.1.1 | BA.2 | BA.2.12.1 | BA.4/5 |
|---|---|---|---|---|---|---|
| Wu01 | | 0.7415 | 0.7126 | 0.7335 | 0.6699 | 0.6575 |
| BA.1 | **0.0065** | | 0.9235 | 0.9608 | 0.8763 | 0.8045 |
| BA.1.1 | **<0.0001** | 0.8001 | | 0.8612 | 0.8861 | 0.7890 |
| BA.2 | **<0.0001** | >0.9999 | >0.9999 | | 0.8661 | 0.7856 |
| BA.2.12.1 | **<0.0001** | **0.0038** | >0.9999 | 0.8001 | | 0.8435 |
| BA.4/5 | **<0.0001** | **<0.0001** | **0.0004** | **0.0002** | 0.1947 | |

*p* value (Dunn's test) — Spearman's Rho

**Extended Data Fig. 1. Serum neutralization of Omicron sublineages.**
**a,** Serum $ID_{50}$s against Wu01 and Omicron sublinages in the cohort of convalescent individuals after infection (V1) and BNT162b2 booster immunization (V2) as in Fig. 2. Lines connect $ID_{50}$s of individual participants at V1 and V2. Dashed lines indicate lower limit of quantification (LLOQ, $ID_{50}$=10). **b,** Numbers in black indicate Spearman's rank correlation coefficients (rho) for comparisons at V2 shown in **c**. Numbers in red indicate *p* values determined by Friedman test followed by Dunn's multiple comparison test for comparisons at V2, with statistically significant differences indicated in bold. **c,** Correlation plots of $log_{10}$ serum $ID_{50}$s against indicated viruses in convalescent individuals at V2. **d,** Serum $ID_{50}$s against Wu01 and Omicron sublineages in the cohort of BNT162b2-vaccinated individuals after the second (V1) and the third vaccine dose (V2) as in Fig. 2. Lines connect $ID_{50}$s of individual participants at V1 and V2. Dashed lines indicate lower limit of quantification (LLOQ, $ID_{50}$=10). **e,** Numbers in black indicate Spearman's rank correlation coefficients (rho) for comparisons at V2 shown in **f**. Numbers in red indicate *p* values determined by Friedman test followed by Dunn's multiple comparison test for comparisons at V2, with statistically significant differences indicated in bold. **f,** Correlation plots of $log_{10}$ serum $ID_{50}$s against indicated viruses in vaccinated individuals at V2. In **a** and **d**, Serum $ID_{50}$s <LLOQ were imputed to ½ LLOQ ($ID_{50}$=5).

### Extended Data Figure 2

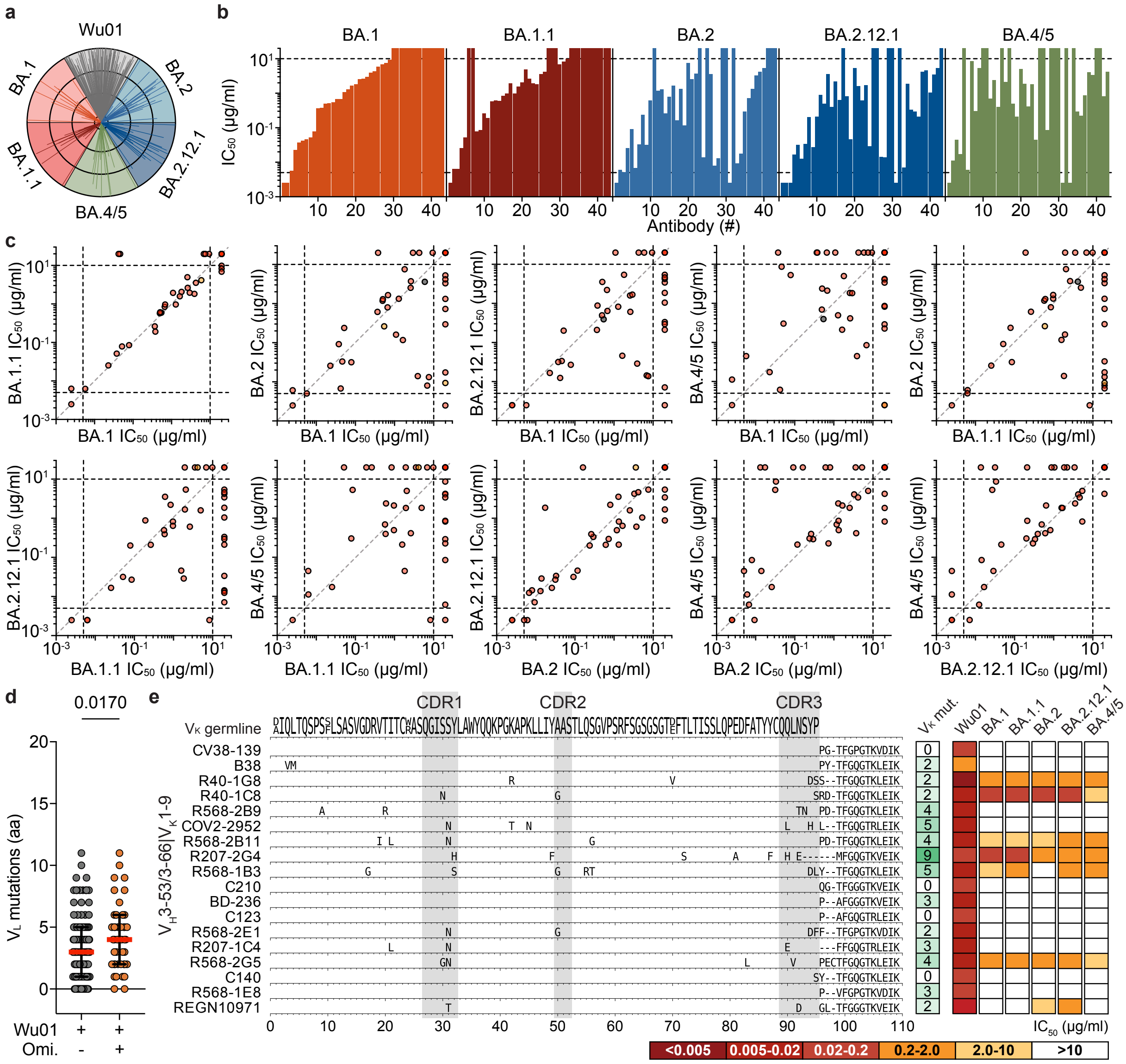

**Extended Data Fig. 2. Neutralization profile of Omicron neutralizing antibodies.**
**a,** Spider plot of $IC_{50}$s for all antibodies against Wu01 and Omicron sublineages. Antibodies are sorted arbitrarily but equally for each virus. Circles indicate $IC_{50}$s (from outer to inner: 0.005, 0.05, 0.5, and 5 µg/ml). **b,** Bar charts of antibodies with neutralizing activity ($IC_{50}$ <10 µg/ml) against any Omicron sublineage (n=43). In each chart, antibodies are sorted by BA.1-neutralizing activity. Dotted lines indicate lower (LLOQ, 0.005 µg/ml) and upper limits of quantification (ULOQ; 10 µg/ml). **c,** $IC_{50}$ correlation plots for antibodies with neutralizing capacity against any Omicron sublineage (n=43). Colors indicate epitopes as in Fig. 3a. Grey dashed lines represent identity lines and black dashed lines indicate limits of quantification. In **a-c**, $IC_{50}$s <LLOQ were imputed to ½ LLOQ ($IC_{50}$=0.0025) and $IC_{50}$s >ULOQ were imputed to 2x ULOQ ($IC_{50}$=20). **d,** Light chain V amino acid mutations of antibodies with neutralizing activity against Wu01 only and antibodies neutralizing both Wu01 and at least one omicron sublineage. Lines indicate medians and interquartile ranges and groups were compared using the two-tailed Mann-Whitney U test. **e,** Light chain sequence alignment of antibodies of the $V_H$3-53/3-66|$V_K$1-9 public clonotype. Letters indicate amino acid (aa) mutations relative to the light chain germline gene. Germline $V_K$ represents the consensus of all identified germline alleles of the depicted antibodies. Number of aa mutations from corresponding germline allele and neutralizing activity against Wu01 and Omicron sublineages are indicated on the right.

### Extended Data Figure 3

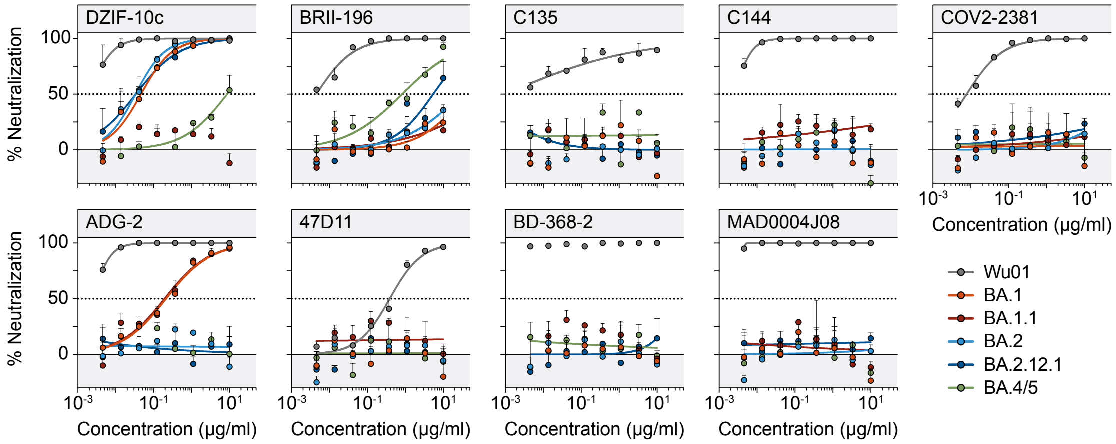


**Extended Data Fig. 3. Omicron sublineage neutralizing activity of monoclonal antibodies in clinical testing.**
Dose response curves showing % neutralization of monoclonal antibodies against Wu01 and Omicron sublineages in a pseudovirus neutralization assay. Circles show averages and error bars indicate standard deviation. Dotted lines indicate 50% neutralization ($IC_{50}$).

#### **Extended Data Table 1** - *Study cohorts*

**a**

| **Convalescent cohort** | |
|---|---|
| **Demographics** | |
| **Participants** - *n* | 20 |
| **Gender** | |
| Female - *n* (%) | 11 (55%) |
| Male - *n* (%) | 9 (45%) |
| **Age** - median years (IQR; range) | 51 (35-60; 25-73) |
| **Reported comorbidities** | |
| Asthma - *n* (%) | 3 (15%) |
| Arterial hypertension - *n* (%) | 2 (10%) |
| Malignancy - *n* (%) | 2 (10%) |
| Gastroesophageal reflux disease - *n* (%) | 1 (5%) |
| **Infection** | |
| **Period of SARS-CoV-2 infection** | February - April 2020 |
| **COVID-19 severity** | |
| Mild symptoms - *n* (%) | 19 (95%) |
| Asymptomatic - *n* (%) | 1 (5%) |
| **Vaccination** | |
| **Vaccine** | BNT162b2 |
| **Time between infection and vaccination** - | |
| median days (IQR; range) | 428 (411-449; 392-483) |
| **Study visits** | |
| **Sampling time point** - median days (IQR; range) | |
| V1 (after disease onset) | 48 (34-58; 22-75) |
| V2 (after vaccination) | 33 (27-52; 23-68) |

**b**

| **Vaccine cohort** | |
|---|---|
| **Demographics** | |
| **Participants** - *n* | 30 |
| **Gender** | |
| Female - *n* (%) | 18 (60%) |
| Male - *n* (%) | 12 (40%) |
| **Age** - median years (IQR; range) | 33 (29-43; 21-59) |
| **Reported comorbidities** | |
| Allergic rhinitis - *n* (%) | 11 (37%) |
| Asthma - *n* (%) | 3 (10%) |
| Gynecologic disease - *n* (%) | 3 (10%) |
| Cardiovascular disease - *n* (%) | 2 (6%) |
| Diabetes - *n* (%) | 2 (6%) |
| Hypothyroidism - *n* (%) | 1 (3%) |
| Chronic liver/Intestinal disease - *n* (%) | 1 (3%) |
| Neurological disorder - *n* (%) | 1 (3%) |
| **Vaccination** | |
| **Vaccine** | BNT162b2 |
| **Time between first and second dose** - | |
| median days (IQR; range) | 21 (21-21; 21-28) |
| **Time between second and third dose** - | |
| median days (IQR; range) | 274 (267-286; 173-307) |
| **Study visits** | |
| **Sampling time point -** median days (IQR; range) | |
| V1 (after second dose) | 28 (27-32; 20-49) |
| V2 (after third dose) | 29 (26-35; 21-57) |

### **Extended Data Table 2** - *Human monoclonal antibody panel analysis*

| # | Name | Epitope | Pseudovirus IC$_{50}$ (μg/ml) | | | | | | Heavy chain | | | | Light chain | | | | Donor | Ref. |
|---|---|---|---|---|---|---|---|---|---|---|---|---|---|---|---|---|---|---|
| | | | Wu01 | BA.1 | BA.1.1 | BA.2 | BA.2.12.1 | BA.4/5 | V gene | GL id. (%)[a] | # aa mut.[b] | CDR3 # aa[c] | V gene | GL id. (%)[a] | # aa mut.[b] | CDR3 # aa[c] | | |
| 1 | **2-7** | RBD | 0.022 | >10 | >10 | 0.296 | 0.333 | 0.292 | 2-5 | 100.0 | 0 | 11 | L2-14 | 93.9 | 6 | 9 | Fab2 | 1 |
| 2 | **2-15** | RBD | <0.005 | >10 | >10 | >10 | >10 | >10 | 1-2 | 95.9 | 4 | 22 | L2-14 | 95.9 | 4 | 10 | Fab2 | 1 |
| 3 | **2-30** | RBD | 0.775 | >10 | >10 | >10 | >10 | >10 | 3-30 | 96.9 | 3 | 12 | K1-9 | 95.8 | 4 | 9 | Fab2 | 1 |
| 4 | **2-36** | RBD | 0.062 | >10 | 9.885 | >10 | >10 | >10 | 4-61 | 96.0 | 4 | 20 | K3-20 | 100.0 | 0 | 9 | Fab2 | 1 |
| 5 | **2-38** | RBD | 2.059 | >10 | >10 | >10 | >10 | >10 | 3-21 | 99.0 | 1 | 14 | L3-19 | 96.8 | 3 | 9 | Fab2 | 1 |
| 6 | **2-43** | RBD | 3.779 | >10 | >10 | >10 | >10 | >10 | 1-2 | 98.0 | 2 | 22 | L2-14 | 96.9 | 3 | 10 | Fab2 | 1 |
| 7 | **4-20** | RBD | 0.029 | >10 | >10 | >10 | >10 | >10 | 1-46 | 95.9 | 4 | 13 | K1-39 | 97.9 | 2 | 10 | Fab4 | 1 |
| 8 | **B38** | RBD | 1.000 | >10 | >10 | >10 | >10 | >10 | 3-53 | 99.0 | 1 | 9 | K1-9 | 97.9 | 2 | 10 | n.a. | 2 |
| 9 | **Bamlanivimab** | RBD | <0.005 | >10 | >10 | >10 | >10 | >10 | 1-69 | 99.0 | 1 | 18 | K1-39 | 97.9 | 2 | 9 | n.a. | 3 |
| 10 | **BD-236** | RBD | 0.018 | >10 | >10 | >10 | >10 | >10 | 3-53 | 96.9 | 3 | 12 | K1-9 | 96.8 | 3 | 9 | Patient 1-64 | 4 |
| 11 | **BD-368-2** | RBD | <0.005 | >10 | >10 | >10 | >10 | >10 | 3-23 | 90.8 | 9 | 18 | K2-28 | 100.0 | 0 | 9 | Patient 1-64 | 4 |
| 12 | **BD23** | RBD | 0.831 | >10 | >10 | >10 | >10 | >10 | 7-4-1 | 100.0 | 0 | 19 | K1-5 | 100.0 | 0 | 9 | Patient 1-64 | 4 |
| 13 | **C002** | RBD | <0.005 | >10 | >10 | >10 | >10 | >10 | 3-30 | 98.0 | 2 | 17 | K1-39 | 98.9 | 1 | 9 | COV21 | 5 |
| 14 | **C022** | RBD | 0.203 | >10 | >10 | >10 | >10 | >10 | 4-39 | 97.0 | 3 | 21 | K1-5 | 97.9 | 2 | 9 | COV21 | 5 |
| 15 | **C101** | RBD | 0.016 | >10 | >10 | >10 | >10 | >10 | 3-53 | 94.8 | 5 | 11 | K3-20 | 96.9 | 3 | 9 | COV107 | 5 |
| 16 | **C102** | RBD | 0.027 | >10 | >10 | >10 | >10 | >10 | 3-53 | 96.9 | 3 | 11 | K3-20 | 100.0 | 0 | 9 | COV107 | 5 |
| 17 | **C104** | RBD | 0.046 | >10 | >10 | >10 | >10 | >10 | 4-34 | 93.8 | 6 | 17 | K3-20 | 94.8 | 5 | 9 | COV107 | 5 |
| 18 | **C105** | RBD | 0.058 | >10 | >10 | >10 | >10 | >10 | 3-53 | 99.0 | 1 | 12 | L2-8 | 98.0 | 2 | 11 | COV107 | 5 |
| 19 | **C123** | RBD | 0.137 | >10 | >10 | >10 | >10 | >10 | 3-53 | 97.9 | 2 | 10 | K1-9 | 100.0 | 0 | 9 | COV107 | 5 |
| 20 | **C125** | RBD | 0.021 | >10 | >10 | >10 | >10 | >10 | 1-58 | 99.0 | 1 | 16 | K3-20 | 100.0 | 0 | 9 | COV107 | 5 |
| 21 | **C128** | RBD | 0.295 | >10 | >10 | >10 | >10 | >10 | 3-23 | 90.7 | 9 | 18 | K3-20 | 93.7 | 6 | 10 | COV072 | 5 |
| 22 | **C135** | RBD | <0.005 | >10 | >10 | >10 | >10 | >10 | 3-30 | 95.9 | 4 | 12 | K1-5 | 96.8 | 3 | 9 | COV072 | 5 |
| 23 | **C140** | RBD | 0.020 | >10 | >10 | >10 | >10 | >10 | 3-66 | 94.8 | 5 | 11 | K1-9 | 100.0 | 0 | 9 | COV072 | 5 |
| 24 | **C144** | RBD | <0.005 | >10 | >10 | >10 | >10 | >10 | 3-53 | 96.9 | 3 | 25 | L2-14 | 99.0 | 1 | 10 | COV047 | 5 |
| 25 | **C155** | RBD | 0.009 | >10 | >10 | >10 | >10 | >10 | 3-53 | 97.9 | 2 | 11 | K3-15 | 98.9 | 1 | 9 | COV047 | 5 |
| 26 | **C165** | RBD | 0.083 | >10 | >10 | >10 | >10 | >10 | 1-69 | 96.9 | 3 | 15 | K3-20 | 99.0 | 1 | 9 | COV072 | 5 |
| 27 | **C210** | RBD | 0.058 | >10 | >10 | >10 | >10 | >10 | 3-53 | 97.9 | 2 | 11 | K1-9 | 100.0 | 0 | 10 | COV96 | 5 |
| 28 | **CC6.31** | RBD | 0.035 | >10 | >10 | >10 | >10 | >10 | 1-46 | 94.9 | 5 | 12 | K1-17 | 100.0 | 0 | 10 | CC6 | 6 |
| 29 | **CC6.33** | RBD | 0.062 | >10 | >10 | >10 | >10 | >10 | 1-69 | 95.9 | 4 | 11 | K3-20 | 98.9 | 1 | 9 | CC6 | 6 |
| 30 | **CC12.4** | RBD | 4.951 | >10 | >10 | >10 | >10 | >10 | 1-2 | 96.9 | 3 | 19 | L2-8 | 96.0 | 4 | 10 | CC12 | 6 |
| 31 | **CnC2t1p1_B4** | RBD | 0.173 | >10 | >10 | >10 | >10 | >10 | 1-18 | 100.0 | 0 | 12 | L2-23 | 99.0 | 1 | 10 | CnC2 | 7 |
| 32 | **CnC2t1p1_D6** | RBD | 1.726 | >10 | >10 | >10 | >10 | >10 | 3-49 | 100.0 | 0 | 17 | K2-28 | 100.0 | 0 | 9 | CnC2 | 7 |
| 33 | **CnC2t1p1_E8** | RBD | 0.449 | >10 | >10 | >10 | >10 | >10 | 1-2 | 85.7 | 14 | 13 | L2-23 | 90.9 | 9 | 10 | CnC2 | 7 |
| 34 | **CnC2t1p1_E12** | RBD | 1.175 | >10 | >10 | >10 | >10 | >10 | 3-49 | 99.0 | 1 | 17 | K2-28 | 100.0 | 0 | 9 | CnC2 | 7 |
| 35 | **CnC2t1p1_G6** | RBD | 5.616 | >10 | >10 | >10 | >10 | >10 | 1-2 | 85.7 | 14 | 13 | L2-23 | 89.9 | 10 | 10 | CnC2 | 7 |
| 36 | **COV2-2050** | RBD | <0.005 | >10 | >10 | >10 | >10 | >10 | 1-2 | 95.9 | 4 | 23 | L1-44 | 94.8 | 5 | 11 | n.a. | 8 |
| 37 | **COV2-2064** | RBD | 0.500 | >10 | >10 | >10 | >10 | >10 | 1-8 | 91.8 | 8 | 16 | L1-44 | 91.8 | 8 | 11 | n.a. | 8 |
| 38 | **COV2-2068** | RBD | 0.045 | 1.283 | 0.968 | 1.330 | 0.821 | 0.497 | 3-53 | 93.8 | 6 | 16 | L1-40 | 96.0 | 4 | 12 | n.a. | 8 |
| 39 | **COV2-2098** | RBD | 0.006 | >10 | >10 | >10 | >10 | >10 | 3-23 | 87.9 | 11 | 11 | K3-15 | 91.6 | 8 | 9 | n.a. | 8 |
| 40 | **COV2-2130** | RBD | <0.005 | 6.848 | >10 | 0.008 | 0.014 | 0.085 | 3-15 | 96.0 | 4 | 22 | K4-1 | 96.0 | 4 | 8 | n.a. | 8 |
| 41 | **COV2-2196** | RBD | <0.005 | >10 | >10 | >10 | 3.438 | >10 | 1-58 | 98.0 | 2 | 16 | K3-20 | 98.9 | 1 | 10 | n.a. | 8 |
| 42 | **COV2-2268** | RBD | 0.510 | >10 | >10 | 3.324 | 4.248 | 4.071 | 2-5 | 98.0 | 2 | 11 | L2-14 | 97.0 | 3 | 11 | n.a. | 8 |
| 43 | **COV2-2308** | RBD | 0.007 | >10 | >10 | >10 | >10 | >10 | 3-23 | 87.9 | 11 | 10 | K3-15 | 91.6 | 8 | 9 | n.a. | 8 |
| 44 | **COV2-2354** | RBD | 0.458 | >10 | >10 | >10 | >10 | >10 | 3-53 | 89.6 | 10 | 12 | L6-57 | 100.0 | 0 | 10 | n.a. | 8 |
| 45 | **COV2-2381** | RBD | 0.008 | >10 | >10 | >10 | >10 | >10 | 1-58 | 98.0 | 2 | 16 | K3-20 | 95.8 | 4 | 10 | n.a. | 8 |
| 46 | **COV2-2479** | RBD | <0.005 | >10 | >10 | >10 | >10 | >10 | 1-69 | 90.8 | 9 | 14 | K3-15 | 96.8 | 3 | 8 | n.a. | 8 |
| 47 | **COV2-2499** | RBD | 0.012 | >10 | >10 | >10 | >10 | >10 | 4-39 | 98.0 | 2 | 19 | L3-19 | 95.8 | 4 | 11 | n.a. | 8 |
| 48 | **COV2-2531** | RBD | 0.366 | >10 | >10 | >10 | >10 | >10 | 4-59 | 90.7 | 9 | 12 | L6-57 | 95.9 | 4 | 9 | n.a. | 8 |
| 49 | **COV2-2539** | RBD | 0.562 | >10 | >10 | >10 | >10 | >10 | 1-8 | 94.9 | 5 | 16 | L1-44 | 92.9 | 7 | 11 | n.a. | 8 |
| 50 | **COV2-2562** | RBD | 0.427 | >10 | >10 | >10 | >10 | >10 | 1-8 | 91.8 | 8 | 16 | L1-44 | 91.8 | 8 | 11 | n.a. | 8 |
| 51 | **COV2-2677** | RBD | 3.904 | >10 | >10 | >10 | >10 | >10 | 4-39 | 99.0 | 1 | 12 | L6-57 | 100.0 | 0 | 10 | n.a. | 8 |
| 52 | **COV2-2678** | RBD | 0.009 | >10 | >10 | >10 | >10 | >10 | 3-20 | 94.8 | 5 | 22 | L3-19 | 96.8 | 3 | 11 | n.a. | 8 |
| 53 | **COV2-2752** | RBD | 0.028 | >10 | >10 | >10 | >10 | 4.205 | 3-53 | 95.9 | 4 | 10 | K1-33 | 97.9 | 2 | 9 | n.a. | 8 |
| 54 | **COV2-2841** | RBD | 0.345 | >10 | >10 | >10 | >10 | >10 | 4-59 | 94.8 | 5 | 12 | L6-57 | 98.0 | 2 | 9 | n.a. | 8 |
| 55 | **COV2-2919** | RBD | 0.071 | >10 | >10 | >10 | >10 | >10 | 2-70 | 97.0 | 3 | 12 | K1-39 | 94.7 | 5 | 9 | n.a. | 8 |
| 56 | **COV2-2952** | RBD | 0.010 | >10 | >10 | >10 | >10 | >10 | 3-66 | 93.8 | 6 | 11 | K1-9 | 94.7 | 5 | 9 | n.a. | 8 |
| 57 | **COV2-2955** | RBD | 0.018 | >10 | >10 | >10 | >10 | >10 | 3-30 | 95.9 | 4 | 22 | K2D-29 | 98.0 | 2 | 9 | n.a. | 8 |
| 58 | **COVA2-29** | RBD | 0.837 | >10 | >10 | >10 | >10 | >10 | 3-30 | 97.9 | 2 | 18 | K1-39 | 98.9 | 1 | 9 | COSCA2 | 9 |
| 59 | **CV-X2-106** | RBD | 0.088 | >10 | >10 | >10 | >10 | >10 | 1-69 | 100.0 | 0 | 18 | K1-39 | 97.9 | 2 | 9 | CV-X2 | 10 |
| 60 | **CV07-262** | RBD | 0.005 | >10 | >10 | >10 | >10 | >10 | 1-2 | 96.9 | 3 | 22 | L2-23 | 98.0 | 2 | 10 | CV07 | 10 |
| 61 | **CV07-270** | RBD | 0.016 | >10 | >10 | >10 | >10 | >10 | 3-11 | 99.0 | 1 | 22 | L2-14 | 98.0 | 2 | 10 | CV07 | 10 |
| 62 | **CV38-139** | RBD | 0.039 | >10 | >10 | >10 | >10 | >10 | 3-66 | 97.9 | 2 | 10 | K1-9 | 100.0 | 0 | 10 | CV38 | 10 |
| 63 | **CV38-142** | RBD | 1.285 | >10 | >10 | >10 | >10 | >10 | 5-51 | 95.9 | 4 | 16 | K1-39 | 100.0 | 0 | 11 | CV38 | 10 |
| 64 | **DH1042** | RBD | 0.008 | >10 | >10 | >10 | >10 | >10 | 1-69 | 96.9 | 3 | 16 | K1-39 | 97.9 | 2 | 9 | Donor 26 | 11 |
| 65 | **DH1128** | RBD | 3.146 | >10 | >10 | >10 | >10 | >10 | 3-23 | 88.8 | 11 | 8 | K6-21 | 94.7 | 5 | 9 | Donor 26 | 11 |
| 66 | **DH1138** | S1* | 2.256 | >10 | >10 | >10 | >10 | >10 | 4-61 | 97.0 | 3 | 12 | K3-11 | 97.9 | 2 | 8 | Donor 26 | 11 |
| 67 | **DH1184** | RBD | 0.007 | >10 | >10 | >10 | >10 | >10 | 1-69 | 95.9 | 4 | 18 | K3-20 | 100.0 | 0 | 9 | Donor 26 | 11 |
| 68 | **DH1210** | RBD | 0.421 | >10 | >10 | >10 | >10 | >10 | 1-69-2 | 91.8 | 8 | 12 | L1-47 | 91.8 | 8 | 12 | Donor 26 | 11 |
| 69 | **DZIF-10c** | RBD | <0.005 | 0.046 | >10 | 0.033 | 0.034 | 8.638 | 3-30 | 88.8 | 11 | 13 | K1-5 | 89.4 | 10 | 9 | HbnC3 | 7 |
| 70 | **Etesevimab** | RBD | 0.018 | >10 | >10 | >10 | >10 | >10 | 3-66 | 96.9 | 3 | 13 | K1-39 | 97.9 | 2 | 11 | n.a. | 12 |
| 71 | **FnC1t2p1_D4** | RBD | 0.008 | >10 | >10 | >10 | >10 | >10 | 7-4-1 | 93.9 | 6 | 11 | K1-33 | 96.8 | 3 | 9 | FnC1 | 7 |
| 72 | **FnC1t2p1_G5** | RBD | 0.012 | >10 | >10 | >10 | >10 | >10 | 7-4-1 | 93.9 | 6 | 11 | K1-33 | 96.8 | 3 | 9 | FnC1 | 7 |
| 73 | **GW01** | RBD | 0.072 | >10 | >10 | >10 | >10 | >10 | 3-43 | 94.9 | 5 | 20 | L1-44 | 100.0 | 0 | 10 | n.a. | 13 |
| 74 | **HbnC2t1p2_D9** | RBD | 0.046 | >10 | >10 | >10 | >10 | >10 | 3-33 | 95.9 | 4 | 19 | K3-11 | 97.9 | 2 | 11 | HbnC2 | 7 |
| 75 | **HbnC3t1p1_C6** | RBD | <0.005 | 9.482 | >10 | >10 | >10 | >10 | 1-58 | 100.0 | 0 | 16 | K3-20 | 99.0 | 1 | 9 | HbnC3 | 7 |
| 76 | **HbnC3t1p1_G4** | RBD | 0.009 | >10 | >10 | >10 | >10 | >10 | 3-66 | 99.0 | 1 | 11 | K3-20 | 97.9 | 1 | 9 | HbnC3 | 7 |
| 77 | **HbnC3t1p2_B10** | RBD | 0.019 | >10 | >10 | >10 | >10 | >10 | 3-66 | 97.9 | 2 | 11 | K3-20 | 95.8 | 3 | 9 | HbnC3 | 7 |
| 78 | **HbnC3t1p2_C6** | RBD | 0.008 | >10 | >10 | >10 | >10 | >10 | 1-58 | 95.9 | 4 | 16 | K3-20 | 99.0 | 1 | 9 | HbnC3 | 7 |
| 79 | **Imdevimab** | RBD | <0.005 | >10 | >10 | 4.173 | 4.624 | 3.392 | 3-30 | 96.9 | 3 | 13 | L2-14 | 93.9 | 6 | 10 | Donor_1 | 14 |
| 80 | **LY-CoV1404** | RBD | <0.005 | <0.005 | <0.005 | <0.005 | <0.005 | <0.005 | 2-5 | 94.9 | 5 | 11 | L2-14 | 91.9 | 8 | 9 | n.a. | 15 |
| 81 | **MAD0004J08** | RBD | <0.005 | >10 | >10 | >10 | >10 | >10 | 1-69 | 95.9 | 4 | 18 | K3-11 | 97.8 | 2 | 5 | n.a. | 16 |
| 82 | **MnC1t3p1_G9** | RBD | 0.592 | >10 | >10 | >10 | >10 | >10 | 3-23 | 89.7 | 10 | 13 | L7-46 | 94.9 | 5 | 9 | MnC1 | 7 |
| 83 | **MnC2t1p1_A3** | RBD | 0.011 | >10 | >10 | >10 | >10 | >10 | 3-66 | 96.9 | 3 | 13 | K1-12 | 100.0 | 0 | 9 | MnC2 | 7 |
| 84 | **MnC2t1p1_C5** | RBD | 0.042 | >10 | >10 | >10 | >10 | >10 | 3-66 | 100.0 | 0 | 13 | K1-12 | 100.0 | 0 | 9 | MnC2 | 7 |
| 85 | **MnC2t2p1_C11** | RBD | 0.031 | >10 | >10 | >10 | >10 | >10 | 1-69 | 95.9 | 4 | 23 | K1-39 | 96.8 | 3 | 9 | MnC2 | 7 |
| 86 | **MnC4t2p1_B3** | RBD | 0.028 | >10 | >10 | >10 | >10 | >10 | 3-9 | 89.9 | 10 | 17 | K1-12 | 94.7 | 5 | 9 | MnC4 | 7 |
| 87 | **MnC4t2p1_D10** | RBD | 5.820 | >10 | >10 | >10 | >10 | >10 | 4-39 | 90.9 | 9 | 20 | K1-17 | 96.8 | 3 | 9 | MnC4 | 7 |
| 88 | **MnC4t2p1_E6** | RBD | 3.971 | >10 | >10 | >10 | >10 | >10 | 3-9 | 91.9 | 8 | 17 | K1-12 | 95.8 | 4 | 9 | MnC4 | 7 |
| 89 | **MnC4t2p2_A4** | RBD | 9.825 | >10 | >10 | >10 | >10 | >10 | 7-4-1 | 91.8 | 8 | 11 | K3-20 | 91.6 | 8 | 9 | MnC4 | 7 |
| 90 | **MnC5t2p1_G1** | RBD | 0.045 | >10 | >10 | >10 | >10 | >10 | 1-58 | 98.0 | 2 | 16 | K3-20 | 96.9 | 3 | 9 | MnC5 | 7 |

*(table continued on next page)*

### **Extended Data Table 2** - *Human monoclonal antibody panel analysis (continued)*

| | | | Pseudovirus IC50 (μg/ml) | | | | | | Heavy chain | | | | Light chain | | | | | |
|---|---|---|---|---|---|---|---|---|---|---|---|---|---|---|---|---|---|---|
| # | Name | Epitope | Wu01 | BA.1 | BA.1.1 | BA.2 | BA.2.12.1 | BA.4/5 | V gene | GL id. (%)[a] | # aa mut.[b] | CDR3 # aa[c] | V gene | GL id. (%)[a] | # aa mut.[b] | CDR3 # aa[c] | Donor | Ref. |
| 91 | **P2B-2F6** | RBD | 0.035 | >10 | >10 | >10 | >10 | >10 | 4-38-2 | 99.0 | 1 | 20 | L2-8 | 100.0 | 0 | 10 | P2 | 17 |
| 92 | **P2C-1F11** | RBD | 0.008 | >10 | >10 | >10 | 5.398 | 0.820 | 3-66 | 95.9 | 4 | 11 | K3-20 | 100.0 | 0 | 8 | P2 | 17 |
| 93 | **R40-1A1** | RBD | 0.007 | >10 | >10 | >10 | >10 | >10 | 1-18 | 83.7 | 16 | 17 | K2-40 | 95.1 | 5 | 9 | R40 | 18 |
| 94 | **R40-1A8** | RBD | <0.005 | >10 | >10 | 0.729 | 0.295 | 0.226 | 3-43 | 93.9 | 6 | 16 | L2-14 | 94.9 | 5 | 10 | R40 | 18 |
| 95 | **R40-1B4** | RBD | <0.005 | >10 | >10 | >10 | >10 | >10 | 1-2 | 90.8 | 9 | 23 | L1-40 | 94.8 | 5 | 9 | R40 | 18 |
| 96 | **R40-1B9** | RBD | 0.012 | >10 | >10 | >10 | >10 | >10 | 2-70 | 96.0 | 4 | 11 | K1-39 | 95.8 | 4 | 9 | R40 | 18 |
| 97 | **R40-1C8** | RBD | 0.012 | 0.079 | 0.085 | 0.032 | 0.027 | 5.324 | 3-53 | 93.8 | 6 | 11 | K1-9 | 97.9 | 2 | 10 | R40 | 18 |
| 98 | **R40-1D3** | RBD | <0.005 | >10 | 8.296 | <0.005 | <0.005 | <0.005 | 3-23 | 88.8 | 11 | 14 | L2-14 | 93.9 | 6 | 10 | R40 | 18 |
| 99 | **R40-1E1** | RBD | 0.214 | >10 | 6.880 | >10 | >10 | >10 | 3-33 | 94.9 | 5 | 24 | K3-20 | 96.9 | 3 | 9 | R40 | 18 |
| 100 | **R40-1E4** | RBD | <0.005 | >10 | >10 | >10 | >10 | >10 | 1-2 | 95.9 | 4 | 13 | K3-20 | 93.7 | 6 | 8 | R40 | 18 |
| 101 | **R40-1G6** | S1* | 0.823 | 0.546 | 0.592 | 0.263 | 0.394 | 0.396 | 3-33 | 89.8 | 10 | 15 | K1-39 | 93.7 | 6 | 9 | R40 | 18 |
| 102 | **R40-1G8** | RBD | <0.005 | 0.495 | 0.575 | 1.178 | 0.497 | 0.692 | 3-53 | 91.8 | 8 | 11 | K1-9 | 97.8 | 2 | 9 | R40 | 18 |
| 103 | **R40-1G12** | RBD | 0.023 | >10 | >10 | >10 | >10 | >10 | 1-2 | 100.0 | 0 | 17 | L2-14 | 99.0 | 1 | 12 | R40 | 18 |
| 104 | **R40-1H4** | RBD | <0.005 | >10 | >10 | >10 | >10 | >10 | 1-2 | 90.8 | 9 | 15 | L2-11 | 97.9 | 2 | 9 | R40 | 18 |
| 105 | **R121-1F1** | RBD | <0.005 | 3.935 | 1.827 | 0.014 | 0.029 | 0.045 | 3-30 | 90.8 | 9 | 14 | L1-40 | 91.9 | 8 | 11 | R121 | 18 |
| 106 | **R121-3F7** | RBD | 0.055 | >10 | >10 | >10 | >10 | >10 | 4-61 | 97.0 | 3 | 19 | L1-40 | 95.9 | 4 | 11 | R121 | 18 |
| 107 | **R121-3F11** | RBD | 0.406 | 0.667 | 0.845 | 0.800 | 2.209 | >10 | 4-59 | 88.7 | 11 | 20 | L1-40 | 96.0 | 4 | 11 | R121 | 18 |
| 108 | **R121-3G2** | RBD | 0.292 | 4.288 | 3.543 | >10 | >10 | >10 | 4-39 | 87.9 | 12 | 21 | K3-20 | 88.5 | 11 | 9 | R121 | 18 |
| 109 | **R200-1B8** | RBD | 0.008 | >10 | >10 | >10 | >10 | >10 | 4-31 | 91.9 | 8 | 19 | L1-40 | 95.9 | 4 | 11 | R200 | 18 |
| 110 | **R200-1B9** | RBD | <0.005 | 0.038 | 0.052 | 0.092 | 0.032 | >10 | 1-58 | 93.9 | 6 | 16 | K3-20 | 94.8 | 5 | 9 | R200 | 18 |
| 111 | **R200-1F9** | RBD | <0.005 | 0.023 | 0.025 | 0.026 | 0.017 | 0.017 | 3-48 | 90.8 | 9 | 15 | K3-11 | 94.7 | 5 | 9 | R200 | 18 |
| 112 | **R200-1G11** | RBD | <0.005 | >10 | >10 | >10 | >10 | >10 | 4-31 | 94.9 | 5 | 18 | K1-39 | 92.6 | 7 | 9 | R200 | 18 |
| 113 | **R200-4F4** | RBD | 0.008 | >10 | >10 | >10 | >10 | >10 | 1-69 | 93.9 | 6 | 25 | K1-33 | 97.9 | 2 | 9 | R200 | 18 |
| 114 | **R207-1C1** | RBD | <0.005 | >10 | >10 | >10 | >10 | >10 | 1-69 | 86.7 | 13 | 14 | K3-15 | 98.9 | 1 | 8 | R207 | 18 |
| 115 | **R207-1C4** | RBD | 0.016 | >10 | >10 | >10 | >10 | >10 | 3-53 | 91.8 | 8 | 11 | K1-9 | 96.8 | 3 | 8 | R207 | 18 |
| 116 | **R207-1G1** | RBD | 0.102 | >10 | >10 | >10 | >10 | >10 | 1-18 | 93.9 | 6 | 12 | L2-23 | 99.0 | 1 | 10 | R207 | 18 |
| 117 | **R207-2A6** | RBD | 0.014 | 2.575 | 4.952 | 2.528 | 1.599 | 1.850 | 3-53 | 89.7 | 10 | 11 | K3-15 | 98.9 | 1 | 9 | R207 | 18 |
| 118 | **R207-2A10** | RBD | <0.005 | >10 | >10 | >10 | >10 | >10 | 1-8 | 92.9 | 7 | 15 | L2-23 | 99.0 | 1 | 11 | R207 | 18 |
| 119 | **R207-2C2** | RBD | 0.010 | >10 | >10 | >10 | >10 | >10 | 3-53 | 91.8 | 8 | 12 | L2-8 | 93.8 | 6 | 10 | R207 | 18 |
| 120 | **R207-2F11** | RBD | <0.005 | <0.005 | 0.006 | 0.006 | <0.005 | 0.011 | 3-53 | 89.7 | 10 | 11 | K1-33 | 90.5 | 9 | 9 | R207 | 18 |
| 121 | **R207-2G4** | RBD | 0.021 | 0.052 | 0.079 | 0.239 | 0.201 | 0.303 | 3-53 | 92.8 | 7 | 11 | K1-9 | 90.4 | 9 | 5 | R207 | 18 |
| 122 | **R207-2H1** | RBD | <0.005 | >10 | >10 | >10 | >10 | >10 | 1-46 | 88.8 | 11 | 18 | L1-40 | 91.7 | 8 | 12 | R207 | 18 |
| 123 | **R259-1B9** | RBD | <0.005 | 0.369 | 0.262 | 0.614 | 0.211 | >10 | 1-58 | 91.8 | 8 | 16 | K3-20 | 94.8 | 5 | 9 | R259 | 18 |
| 124 | **R339-1B11** | RBD | <0.005 | 1.090 | 1.946 | 0.161 | >10 | >10 | 3-7 | 88.8 | 11 | 16 | K2-28 | 97.0 | 3 | 9 | R339 | 18 |
| 125 | **R339-3B5** | S2 | 1.158 | 5.954 | 4.139 | 3.596 | >10 | >10 | 1-46 | 81.6 | 18 | 11 | K3-20 | 92.7 | 7 | 11 | R339 | 18 |
| 126 | **R339-3C6** | S1* | 2.657 | >10 | >10 | >10 | >10 | >10 | 4-59 | 92.8 | 7 | 20 | K2D-29 | 89.0 | 11 | 9 | R339 | 18 |
| 127 | **R410-1A8** | RBD | 0.347 | 0.507 | 0.610 | 1.304 | 3.547 | 2.394 | 2-5 | 92.9 | 7 | 15 | K1D-12 | 94.7 | 5 | 9 | R410 | 18 |
| 128 | **R568-1A9** | RBD | <0.005 | 7.558 | >10 | 0.013 | 0.014 | >10 | 1-69 | 88.8 | 11 | 17 | K1-5 | 94.7 | 5 | 10 | R568 | 18 |
| 129 | **R568-1B3** | RBD | 0.010 | 2.954 | 1.954 | >10 | 1.678 | 1.829 | 3-53 | 91.8 | 8 | 11 | K1-9 | 94.6 | 5 | 9 | R568 | 18 |
| 130 | **R568-1C6** | RBD | <0.005 | >10 | >10 | 0.017 | 1.862 | >10 | 1-69 | 92.8 | 7 | 18 | K1-5 | 97.9 | 2 | 10 | R568 | 18 |
| 131 | **R568-1E8** | RBD | 0.048 | >10 | >10 | >10 | >10 | >10 | 3-53 | 96.9 | 3 | 11 | K1-9 | 96.8 | 3 | 9 | R568 | 18 |
| 132 | **R568-1G9** | RBD | 0.007 | 0.006 | 0.006 | 0.005 | <0.005 | 0.045 | 3-66 | 96.9 | 3 | 12 | L1-40 | 99.0 | 1 | 10 | R568 | 18 |
| 133 | **R568-2A1** | RBD | 0.295 | >10 | >10 | >10 | >10 | >10 | 3-11 | 95.9 | 4 | 15 | L1-47 | 99.0 | 1 | 11 | R568 | 18 |
| 134 | **R568-2A3** | RBD | <0.005 | >10 | >10 | >10 | >10 | >10 | 1-2 | 89.8 | 10 | 15 | L2-8 | 97.0 | 3 | 10 | R568 | 18 |
| 135 | **R568-2B9** | RBD | 0.014 | >10 | >10 | >10 | >10 | >10 | 3-53 | 89.7 | 10 | 11 | K1-9 | 95.8 | 4 | 10 | R568 | 18 |
| 136 | **R568-2B11** | RBD | 0.014 | 2.638 | 2.624 | 3.749 | 0.609 | 0.416 | 3-53 | 93.8 | 6 | 11 | K1-9 | 95.8 | 4 | 10 | R568 | 18 |
| 137 | **R568-2E1** | RBD | 0.006 | >10 | >10 | >10 | >10 | >10 | 3-53 | 93.8 | 6 | 11 | K1-9 | 97.8 | 2 | 9 | R568 | 18 |
| 138 | **R568-2E7** | S1* | 0.007 | >10 | >10 | 0.009 | 0.007 | <0.005 | 3-23 | 90.7 | 9 | 8 | K3-15 | 97.9 | 2 | 10 | R568 | 18 |
| 139 | **R568-2F1** | RBD | 0.009 | >10 | >10 | >10 | >10 | >10 | 1-69 | 89.7 | 10 | 17 | L1-47 | 94.8 | 5 | 12 | R568 | 18 |
| 140 | **R568-2G5** | RBD | 0.006 | 0.684 | 0.973 | 1.648 | 0.640 | 2.122 | 3-53 | 91.8 | 8 | 11 | K1-9 | 95.8 | 4 | 11 | R568 | 18 |
| 141 | **R568-2G11** | RBD | <0.005 | 0.042 | >10 | 0.007 | 0.012 | 0.006 | 3-15 | 92.0 | 8 | 15 | K1-8 | 95.8 | 4 | 9 | R568 | 18 |
| 142 | **R616-1A11** | RBD | 0.037 | >10 | >10 | >10 | >10 | >10 | 1-69 | 88.7 | 11 | 16 | K1-5 | 93.7 | 6 | 11 | R616 | 18 |
| 143 | **R616-1D6** | RBD | 3.278 | >10 | >10 | >10 | >10 | >10 | 4-34 | 83.5 | 16 | 20 | K1-39 | 90.5 | 9 | 9 | R616 | 18 |
| 144 | **R616-1F10** | RBD | 0.052 | >10 | >10 | >10 | >10 | >10 | 3-30 | 86.7 | 13 | 17 | L1-44 | 94.9 | 5 | 12 | R616 | 18 |
| 145 | **R616-1G4** | RBD | 0.005 | >10 | >10 | >10 | >10 | >10 | 3-53 | 96.9 | 3 | 15 | L2-14 | 97.0 | 3 | 12 | R616 | 18 |
| 146 | **R849-1C11** | RBD | 0.409 | 1.811 | 2.062 | 7.581 | 5.386 | 5.023 | 4-30-4 | 94.9 | 5 | 27 | L2-18 | 97.0 | 3 | 10 | R849 | 18 |
| 147 | **R849-1G7** | RBD | <0.005 | >10 | >10 | >10 | >10 | >10 | 1-2 | 90.8 | 9 | 16 | L2-23 | 91.8 | 8 | 10 | R849 | 18 |
| 148 | **R849-1H1** | RBD | 0.132 | >10 | >10 | >10 | >10 | >10 | 3-33 | 96.9 | 3 | 20 | K1-39 | 96.8 | 3 | 10 | R849 | 18 |
| 149 | **R849-3H2** | RBD | 0.013 | >10 | >10 | >10 | >10 | >10 | 3-30 | 92.9 | 7 | 14 | K1-39 | 97.9 | 2 | 8 | R849 | 18 |
| 150 | **REGN10954** | RBD | 0.009 | 1.641 | 1.583 | 0.117 | 0.046 | 0.212 | 3-66 | 96.9 | 3 | 13 | K1-33 | 94.7 | 5 | 9 | Donor_3 | 14 |
| 151 | **REGN10955** | RBD | 0.017 | >10 | >10 | 1.319 | 0.212 | 0.857 | 3-66 | 95.9 | 4 | 9 | K1-33 | 97.9 | 2 | 9 | Donor_3 | 14 |
| 152 | **REGN10964** | RBD | <0.005 | >10 | >10 | >10 | >10 | >10 | 4-59 | 94.8 | 5 | 12 | K1-39 | 93.7 | 6 | 9 | Donor_1 | 14 |
| 153 | **REGN10970** | RBD | 0.014 | >10 | >10 | >10 | >10 | >10 | 3-66 | 96.9 | 3 | 14 | K1-33 | 98.9 | 1 | 9 | Donor_1 | 14 |
| 154 | **REGN10971** | RBD | 0.008 | >10 | >10 | 5.290 | 1.046 | >10 | 3-53 | 95.9 | 4 | 11 | K1-9 | 97.9 | 2 | 10 | Donor_1 | 14 |
| 155 | **REGN10977** | RBD | <0.005 | >10 | >10 | >10 | >10 | >10 | 1-69 | 95.9 | 4 | 16 | K3-20 | 95.8 | 4 | 9 | Donor_1 | 14 |
| 156 | **REGN10986** | RBD | 0.006 | 0.381 | 0.192 | >10 | 0.885 | >10 | 3-66 | 97.9 | 2 | 13 | L1-40 | 98.0 | 2 | 12 | Donor_1 | 14 |
| 157 | **REGN10989** | RBD | <0.005 | >10 | >10 | >10 | >10 | >10 | 1-2 | 93.9 | 6 | 16 | L2-14 | 92.9 | 7 | 10 | Donor_3 | 14 |
| 158 | **S2X35** | RBD | 0.057 | >10 | >10 | >10 | >10 | >10 | 1-18 | 98.0 | 2 | 21 | L1-40 | 99.0 | 1 | 13 | Donor S2X | 19 |

a Amino acid identity relative to germline gene (framework region 1 to framework region 3).
b Number of amino acid mutations relative to germline gene (framework region 1 to framework region 3).
c Length of CDR3 in amino acids.
* Indicates epitope in S1 domain of SARS-CoV-2 spike protein outside of receptor-binding domain (e.g., N-terminal domain).
n.a. Indicates that donor ID was not unambiguously identified.
RBD Receptor-binding domain.
Ref. References:

1, Liu L. et al., 2020; 2, Wu Y. et al., 2020; 3, Jones B.E. et al., 2021; 4, Cao Y. et al., 2020; 5, Robbiani D.F. et al., 2020; 6, Rogers T.F. et al., 2020; 7, Kreer C. et al, 2020; 8, Zost S.J. et al, 2020; 9, Brouwer P.J.M. et al., 2020; 10, Kreye J. et al, 2020; 11, Li D. et al, 2021; 12, Shi R. et al., 2020; 13, Wang Y. et al, 2022; 14, Hansen J. et al., 2020; 15, Westendorf K. et al, 2022; 16, Andreano E. et al., 2021; 17, Ju B. et al., 2020; 18, Vanshylla K. et al., 2022; 19, Piccoli L. et al, 2020.
